# NF1 regulates mesenchymal glioblastoma plasticity and aggressiveness through the AP-1 transcription factor FOSL1

**DOI:** 10.1101/834531

**Authors:** Carolina Marques, Thomas Unterkircher, Paula Kroon, Annalisa Izzo, Yuliia Dramaretska, Eva Kling, Barbara Oldrini, Oliver Schnell, Sven Nelander, Erwin F. Wagner, Latifa Bakiri, Gaetano Gargiulo, Maria Stella Carro, Massimo Squatrito

## Abstract

The molecular basis underlying Glioblastoma (GBM) heterogeneity and plasticity are not fully understood. Using transcriptomic data of patient-derived brain tumor stem cell lines (BTSCs), classified based on GBM-intrinsic signatures, we identify the AP-1 transcription factor *FOSL1* as a key regulator of the mesenchymal (MES) subtype. We provide a mechanistic basis to the role of the Neurofibromatosis type 1 gene (*NF1*), a negative regulator of the RAS/MAPK pathway, in GBM mesenchymal transformation through the modulation of *FOSL1* expression. Depletion of *FOSL1* in *NF1*-mutant human BTSCs and *Kras*-mutant mouse neural stem cells results in loss of the mesenchymal gene signature, reduction in stem cell properties and *in vivo* tumorigenic potential. Our data demonstrate that *FOSL1* controls GBM plasticity and aggressiveness in response to *NF1* alterations.

## Introduction

Gliomas are the most common primary brain tumor in adults. Given the strong association of the *isocitrate dehydrogenase 1* and *2* (*IDH1/2*) genes mutations with glioma patients survival, the 2016 WHO classification, that integrates both histological and molecular features, has introduced the distinction of *IDH*-wildtype (IDH-wt) or *IDH*-mutant (IDH-mut) in diffuse gliomas ^1^. IDH-wt glioblastoma (GBM) represents the most frequent and aggressive form of gliomas, characterized by high molecular and cellular inter- and intra-tumoral heterogeneity.

Large-scale sequencing approaches have evidenced how concurrent perturbations of cell cycle regulators, growth and survival pathways, mediated by RAS/MAPK and PI3K/AKT signaling, play a significant role in driving adult GBMs ^2–4^. Moreover, various studies have classified GBM in different subtypes, using transcriptional profiling, being now the Proneural (PN), Classical (CL) and Mesenchymal (MES) the most widely accepted ^3,5,6^. Patients with the MES subtype tend to have worse survival rates compared to other subtypes, both in the primary and recurrent tumor settings (Wang et al., 2017). The most frequent genetic alterations – Neurofibromatosis type 1 gene (*NF1*) copy number loss or mutation – and important regulators of the MES subtype, such as *STAT3*, *CEBPB* and *TAZ*, have been identified^3,7,8^. Nevertheless, the mechanisms of regulation of MES GBMs are still not fully understood. For example, whether the MES transcriptional signature is controlled through tumor cell-intrinsic mechanisms or influenced by the tumor microenvironment (TME) is still an unsolved question. In fact, the critical contribution of the TME adds another layer of complexity to MES GBMs. Tumors from this subtype are highly infiltrated by non-neoplastic cells, as compared to PN and CL subtypes ^6^. Additionally, MES tumors express high levels of angiogenic markers and exhibit high levels of necrosis ^9^.

Even though each subtype is associated with specific genetic alterations, there is a considerable plasticity among them: different subtypes co-exist in the same tumors and shifts in subtypes can occur over time ^10,11^. This plasticity may be explained by acquisition of new genetic and epigenetic abnormalities, by stem-like reprogramming or by clonal variation ^12^. It is also not fully understood whether the distinct subtypes evolve from a common glioma precursor ^13^. For instance, PN tumors often switch phenotype to MES upon recurrence, and treatment also increases the mesenchymal gene signature, suggesting that MES transition, or epithelial to mesenchymal (EMT)-like, in GBM is associated with tumor progression and therapy resistance ^5,14,15^. Yet, the frequency and relevance of this EMT-like phenomenon in glioma progression remains unclear. EMT has also been associated with stemness in other cancers ^16–18^. Glioma stem cells (GSCs) share features with normal neural stem cells (NSCs) such as self-renewal and ability to differentiate into distinct cellular lineages (astrocytes, oligodendrocytes and neurons) but are thought to be the responsible for tumor relapse, given their ability to repopulate tumors and their resistance to treatment ^19,20^.

*FOSL1*, that encodes FRA-1, is an AP-1 transcription factor with prognostic value in different epithelial tumors, where its overexpression correlates with tumor progression or worse patient survival ^21–26^. Moreover, the role of *FOSL1* in EMT has been documented in breast and colorectal cancers ^27–29^. In GBM, it has been shown that *FOSL1* modulates *in vitro* glioma cell malignancy ^30^.

Here we report that *NF1* loss, by increasing RAS/MAPK activity, modulates *FOSL1* expression which in turn plays a central function in the regulation of MES GBM. Using a surrogate mouse model of MES GBM and patient-derived MES brain tumor stem cells (BTSCs), we show that *FOSL1* is responsible for sustaining cell growth *in vitro* and *in vivo*, and for the maintenance of stem-like properties. We propose that *FOSL1* is an important regulator of GBM stemness, MES features and plasticity, controlling an EMT-like process with therapeutically relevant implications.

## Results

### *FOSL1* is a key regulator of the MES subtype

To study the tumor cell-intrinsic signaling pathways that modulate the GBM expression subtypes we assembled a collection of transcriptomic data (both expression arrays and RNA-sequencing) of 144 samples derived from 116 independent BTSC lines (see Methods for details). Samples were then classified according to the previously reported 50-gene glioma-intrinsic transcriptional subtype signatures and the single sample gene set enrichment analysis (ssGSEA)-based equivalent distribution resampling classification strategy ^6^. Overall, 40% of the samples were identified as CL, 38% as MES and 22% as PN (Supplementary Table 1). Principal component analysis showed a large overlap of the transcription profile among CL and PN BTSCs while most of the MES BTSCs appeared as a separate group (Fig. 1a). Differential gene expression analysis comparing MES versus Non-MES (PN and CL) BTSCs confirmed a clear separation among the two groups, with the exception of a small number of cell lines that showed a mixed expression profile (Fig. 1b and Supplementary Table 2).

**Fig. 1.**
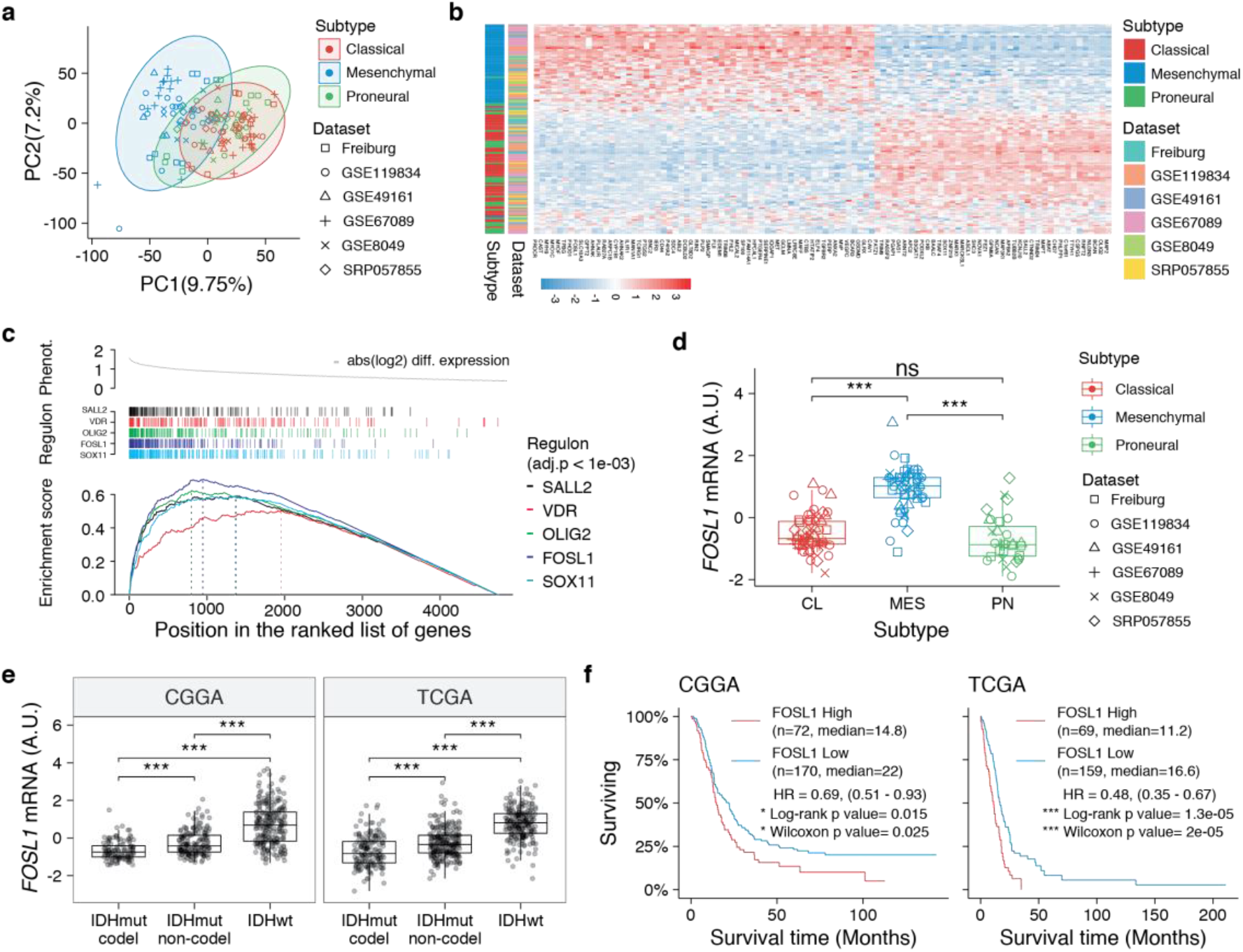
*FOSL1* is a *bona fide* regulator of the glioma-intrinsic MES transcriptional signature. **a,** Principal Component (PC) analysis of the BTSCs expression dataset. **b,** Heatmap of the top 100 differentially expressed genes between MES and Non-MES BTSCs. **c,** One-tail GSEA of the top 5 scoring TFs in the MRA. **d,** *FOSL1* mRNA expression in the BTSCs dataset. One-way ANOVA with Tukey multiple pairwise-comparison, ***P ≤ 0.001, ns = not-significant. **e,** *FOSL1* mRNA expression in the CGGA and TCGA datasets. Tumors were separated according to their molecular subtype classification. One-way ANOVA with Tukey multiple pairwise-comparison, ***P ≤ 0.001. **f,** Kaplan-Meier survival curves of IDH-wt gliomas in the CGGA and TCGA datasets stratified based on *FOSL1* expression.

To reveal the signaling pathways underlying the differences between MES and Non-MES BTSCs we then applied a network-based approach based on ARACNe (Algorithm for the Reconstruction of Accurate Cellular Networks) ^8,31^, which identifies a list of transcription factors (TFs) with their predicted targets, defined as regulons. The regulon for each TF is constituted by all the genes whose expression data exhibit significant mutual information with that of a given TF and are thus expected to be regulated by that TF ^32,33^. Enrichment of a relevant gene signature in each of the regulons can point to the TFs acting as master regulators (MRs) of the response or phenotype ^8,33^. Master regulator analysis (MRA), identified a series of TFs, among which *SOX11*, *FOSL1*, *OLIG2*, *VDR* and *SALL2* were the top 5 most statistically significant (Benjamini-Hochberg P < 0.0001) (Fig. 1c and Supplementary Table 3). *FOSL1* and *VDR* were significantly upregulated in the MES BTSCs, while *SOX11*, *OLIG2* and *SALL2* were upregulated in the Non-MES BTSCs (Fig. 1d and Extended Data Fig. 1a). Gene set enrichment analysis (GSEA) evidenced how the regulons for the top 5 TFs are enriched for genes that are differentially expressed among the two classes (MES and Non-MES) with *FOSL1* having the highest enrichment score (Fig. 1c and Extended Data Fig. 1b).

We then analyzed the CGGA and TCGA pan-glioma datasets ^34,35^ and observed that *FOSL1* expression is elevated in the IDH-wt glioma molecular sub-group (Fig. 1e and Supplementary Table 4) with a significant upregulation in the MES subtype both in bulk tumors and also at the single-cell level (Extended Data Fig. 1c,d). Importantly, high expression levels were associated with worse prognosis in IDH-wt tumors (Fig. 1f), thus suggesting that *FOSL1* could represent not only a key regulator of the glioma-intrinsic MES signature, but also a putative key player in MES glioma pathogenesis.

### *NF1* modulates the MES signature and *FOSL1* expression

*NF1* alterations and activation of the RAS/MAPK signaling have been previously associated with the MES GBM subtype ^2,3,6,36^. However, whether *NF1* plays a broader functional role in the regulation of the MES gene signature (MGS) in IDH-wt gliomas still remains to be established.

We initially grouped, according to the previously described GBM subtype-specific gene signatures, a subset of IDH-wt glioma samples of the TCGA dataset for which RNA-seq data were available (n = 229) (see methods for details). By analyzing the frequency of *NF1* alterations (either point mutations or biallelic gene loss) we confirmed a significant enrichment of *NF1* alterations in MES versus Non-MES tumors (Fisher’s Exact test P = 0.03) (Fig. 2a,b). Importantly, we detected higher level of *FOSL1* mRNA in the cohort of IDH-wt gliomas with *NF1* alterations (Student’s t test P = 0.0018) (Fig. 2c), as well as a significant negative correlation between *FOSL1* and *NF1* mRNA levels (Pearson R = −0.44, P = 7.8e-12) (Fig. 2d and Supplementary Table 4).

**Fig. 2.**
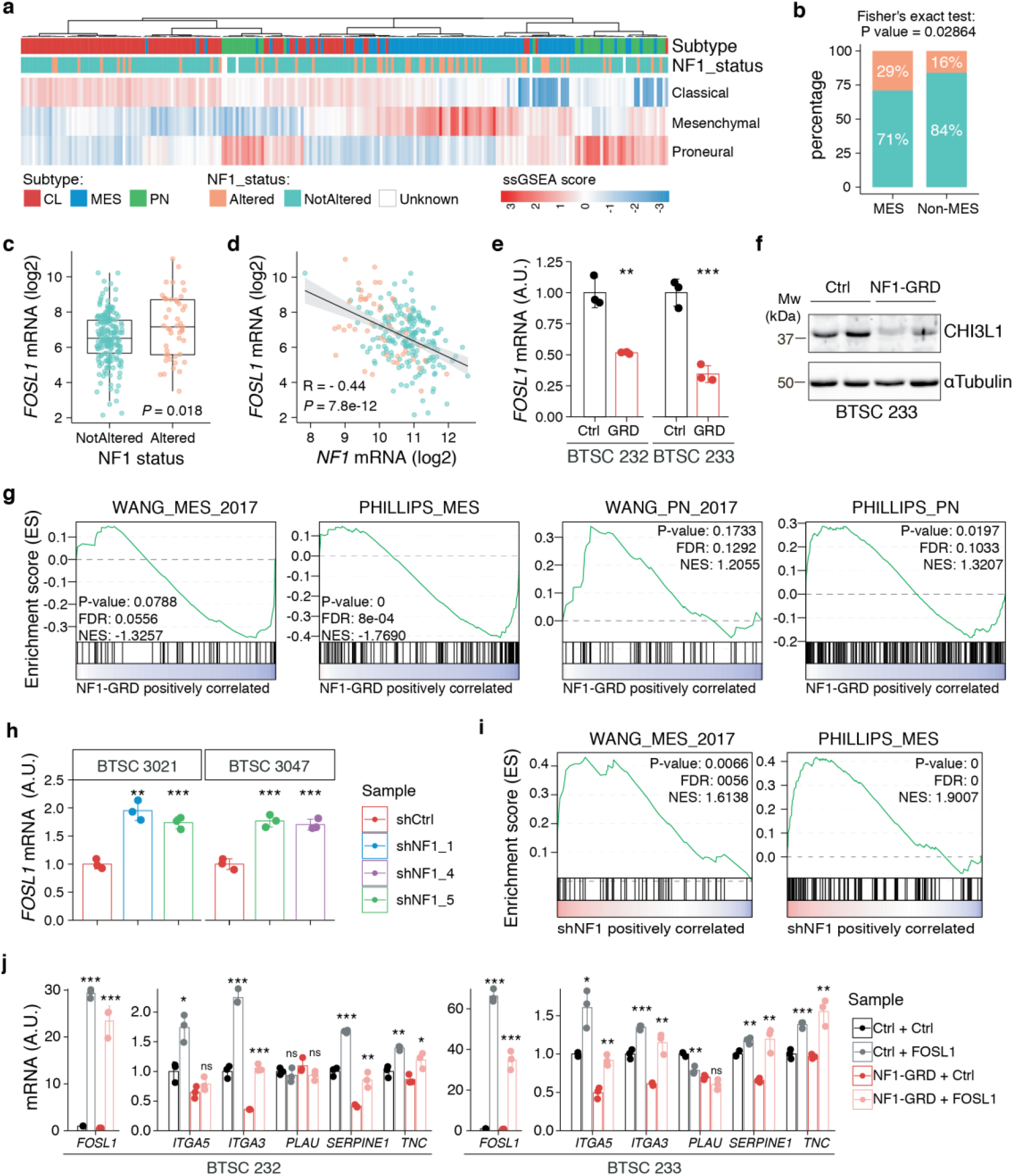
*NF1* is a functional modulator of MES transcriptional signature through *FOSL1* expression regulation. **a,** Heatmap of the subtype ssGSEA scores and *NF1* genetic alterations of the IDH-wt gliomas in the TCGA dataset. **b,** Frequency of *NF1* alterations in MES and Non-MES IDH-wt gliomas. Colors are as in panel a. **c,** *FOSL1* mRNA expression in IDH-wt gliomas, stratified according to *NF1* alterations. Colors are as in panel A. Student’s t test, P = 0.018. **d,** Correlation of *FOSL1* and *NF1* mRNA expression in IDH-wt gliomas. Colors are as in panel (**a**). Pearson correlation, R = −0.044, P = 7.8e-12. **e,** qRT-PCR analysis of *FOSL1* expression upon NF1-GRD overexpression in BTSC 232 and BTSC 233 cells. **f,** Western blot analysis of whole-cell-extract of BTSC 233 cells showing CHI3L1 mesenchymal marker expression upon NF1-GRD transduction; α-Tubulin was used as loading control. **g,** GSEA of BTSC 233 cells transduced with NF1-GRD expressing lentivirus versus Ctrl. Gene signatures from Wang and Phillips studies were analyzed (MES, *left panels*; PN, *right panels*). NES = Normalised enrichment score. **h,** qRT-PCR analysis of *FOSL1* expression upon *NF1* knockdown in BTSC 3021 and BTSC 3047 cells. **i)** GSEA of BTSC 3021 transduced with sh*NF1*_5 versus Ctrl. **j,** qRT-PCR analysis of MES genes expression upon NF1-GRD and *FOSL1* co-expression in BTSC 232 and BTSC 233 cells. qRT-PCR data in (**e**) and (**j**) are presented as mean ± SD (n=3), normalized to 18s rRNA expression; Student’s t test, *P ≤ 0.05, **P ≤ 0.01, ***P ≤ 0.001, ns = not-significant.

To test whether a NF1-MAPK signaling is involved in the regulation of *FOSL1* and the MES subtype, we manipulated *NF1* expression in patient derived GBM tumorspheres of either MES or Non-MES subtypes. To recapitulate the activity of the full-length NF1 protein we transduced the cells with the NF1 GTPase-activating domain (NF1-GRD), spanning the whole predicted Ras GTPase-activating (GAP) domain ^37^. NF1-GRD expression in the MES cell line BTSC 233 led to inhibition of RAS activity as confirmed by analysis of pERK expression upon EGF or serum stimulation (Extended Data Fig. 2a,b) as well as by RAS pull down assay (Extended Data Fig. 2c). Furthermore, analysis of a RAS-induced oncogenic signature expression by GSEA showed a strong reduction in NF1-GRD expressing cells (NES = −1.7; FDR q-value < 0.001) (Extended Data Fig. 2d). Consistent with the negative correlation of *FOSL1* and *NF1* mRNA levels in IDH-wt gliomas (Fig. 2d), NF1-GRD overexpression in two independent MES GBM lines (BTSC 233 and BTSC 232) was associated with a significative downregulation of *FOSL1* and *FOSL1*-regulated genes (Fig. 2e and Extended Data Fig. 3a-c). Concurrently, we also observed a significant decrease of two well characterized mesenchymal features, namely CHI3L1 expression (Fig. 2f) as well as the ability of MES GBM cells to differentiate into osteocytes, a feature shared with mesenchymal stem cells ^38,39^ (Extended Data Fig. 3d). Moreover, NF1-GRD expression led to a significant reduction of the MGSs (Wang signature: NES = −1.3; FDR q-value = 0.05; Phillips signature: NES = −1.7; FDR q-value < 0.001) (Fig. 2g, *left panels*) and to an upregulation of Proneural gene signatures (PNGSs) (Wang signature: NES = 1.2; FDR q-value = 0.12; Phillips signature: NES = 1.3; FDR q-value = 0.1) (Fig. 2g, *right panels*).

Conversely, *NF1* knockdown with three independent shRNAs (shNF1_1, shNF1_4 and shNF1_5), in two Non-MES lines (BTSC 3021 and BTSC 3047) (Extended Data Fig. 3e) led to an upregulation of *FOSL1* (Fig. 2h) with a concomitant significant increase in its targets (Extended Data Fig. 3f,g) and an upregulation of the MGSs (Wang: NES = 1.61; FDR q-value = 0.005; Phillips: NES = 1.9; FDR q-value < 0.001) (Fig. 2i). The PNGSs instead were not significantly lost (data not shown).

The observed NF1-mediated gene expression changes might be potentially driven by an effect on *FOSL1* or other previously described mesenchymal TFs (such as *BHLHB2*, *CEBPB*, *FOSL2*, *RUNX1*, *STAT3* and *TAZ*) ^7,8^. Interestingly, only *FOSL1,* and to some extent *CEBPB,* was consistently downregulated upon NF1-GRD expression (Extended Data Fig. 3h) and upregulated following *NF1* knockdown (Extended Data Fig. 3i). To then test whether *FOSL1* was playing a direct role in the *NF1*-mediated regulation of mesenchymal genes expression, we overexpressed *FOSL1* in the MES GBM lines transduced with the NF1-GRD (Extended Data Fig. 3j). qRT-PCR analysis showed that *FOSL1* was able to rescue the NF1-GRD-mediated downregulation of mesenchymal genes, such as *ITGA3, ITGA5, SERPINE1* and *TNC*. Lastly, exposure of *NF1* silenced cells to the MEK inhibitor GDC-0623, led to a reduction of *FOSL1* upregulation, both at the protein and the mRNA levels, as well as to a downregulation of the mesenchymal genes *ITGA3* and *SERPINE1* (Extended Data Fig. 4a,b).

Overall these evidences suggest that a NF1-MAPK signaling is directly involved in the regulation of the MGS, very likely through the modulation of *FOSL1* expression.

### *Fosl1* deletion induces a shift from a MES to a PN gene signature

To further explore the NF1-MAPK-FOSL1 axis in MES GBM we used a combination of the RCAS-Tva system with the CRISPR/Cas9 technology, recently developed in our laboratory (ref ^40^) to induce *Nf1* loss or *Kras* mutation. Mouse neural stem cells (NSCs) from *hGFAP-Tva; hGFAP-Cre; Trp53^lox^; ROSA26-LSL-Cas9* pups were isolated and infected with viruses produced by DF1 packaging cells transduced with RCAS vectors targeting the expression of *Nf1* through shRNA and sgRNA (sh*Nf1* and sg*Nf1*) or overexpressing a mutant form of *Kras* (*Kras^G12V^*). Loss of NF1 expression was confirmed by western blot and FRA-1 was upregulated in the two models of *Nf1* loss compared to parental cells, and further upregulated in cells infected with *Kras^G12V^* (Fig. 3a). Consistent with activation of the Ras signaling, as result of both *Nf1* loss and *Kras* mutation, the MEK/ERK pathway was more active in infected cells compared to parental cells (Fig. 3a). Higher levels of activation of the MEK/ERK pathway were associated with the induction of mesenchymal genes such as *Plau*, *Plaur*, *Timp1* and *Cd44* (Fig. 3b). Moreover, the upregulation of both FRA-1 and the mesenchymal genes, was blocked by exposing sh*Nf1* and *Kras* mutant cells to the MAPK inhibitors Trametinib or U0126 (Extended Data Fig. 4c,d). These data indicated that *Kras^G12V^*– transduced cells are a suitable model to functionally study the role of a MAPK-FOSL1 axis in MES GBM.

**Fig. 3.**
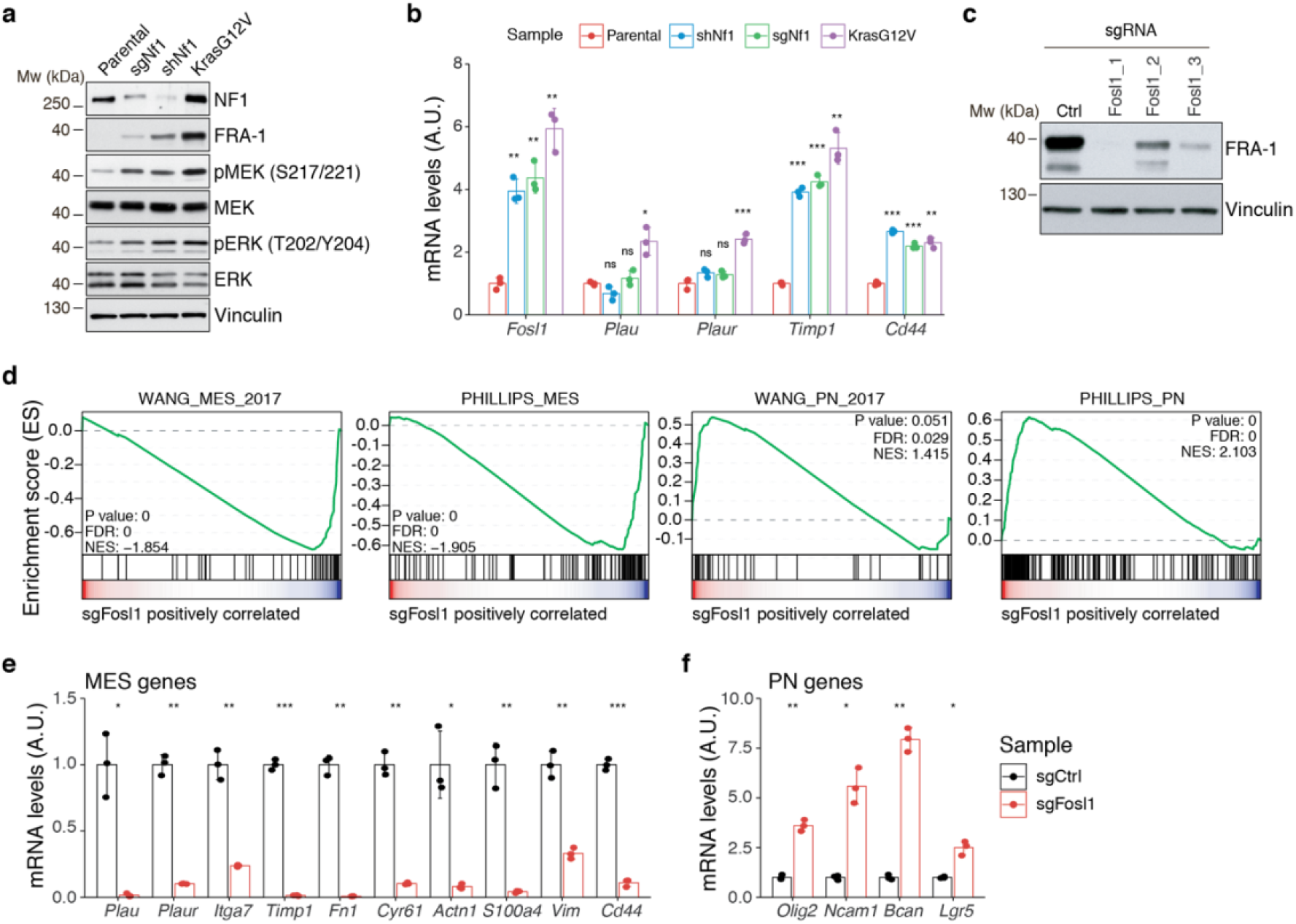
*Fosl1* is induced by MAPK kinase activation and is required for MES gene expression. **a,** Western blot analysis using the specified antibodies of p53-null NSCs, parental and infected with sg*Nf1*, sh*Nf1* and *Kras^G12V^*; Vinculin used as loading control. **b,** mRNA expression of *Fosl1* and MES genes (*Plau, Plaur, Timp1* and *Cd44*), in infected p53-null NSCs, compared to parental cells (not infected). Data from a representative of two experiments are presented as mean ± SD (n=3), normalized to *Gapdh* expression. Student’s t test, relative to parental cells: ns = not significant, *P ≤ 0.05, **P ≤ 0.01, ***P ≤ 0.001. **c,** FRA-1 expression detected by Western blot in p53-null *Kras^G12V^* NSCs upon transduction with sgRNAs targeting *Fosl1*, after selection with 1 μg/mL puromycin; Vinculin used as loading control. **d,** GSEA of p53-null *Kras^G12V^* sg*Fosl1*_1 versus sgCtrl NSCs. Gene signatures from Wang and Phillips studies were analyzed (MES, *left panels*; PN, *right panels*). **e,f,** mRNA expression of MES (**e**) and PN genes (**f**), in sgCtrl and sg*Fosl1*_1 p53-null *Kras^G12V^* NSCs. Data from a representative of two experiments are presented as mean ± SD (n=3), normalized to *Gapdh* expression. Student’s t test, relative to sgCtrl: *P ≤ 0.05; **P ≤ 0.01; ***P ≤ 0.001.

Taking advantage of the Cas9 expression in the generated cell p53-null *Kras^G12V^* NSCs model, *Fosl1* was knocked out through sgRNAs. Efficient downregulation of FRA-1 was achieved with 2 different sgRNAs (Fig. 3c). Cells transduced with sg*Fosl1*_1 and sg*Fosl1*_3 were then subjected to further studies.

As suggested by the data presented here on the human BTSCs datasets and cell lines, *FOSL1* appears to be a key regulator of the MES subtype. Consistently, RNA-seq analysis followed by GSEA of p53-null *Kras^G12V^* sg*Fosl1*_1 versus sgCtrl revealed a significant loss of Wang and Phillips MGSs (Wang: NES = −1.85; FDR q-value < 0.001; Phillips: NES = −1.91; FDR q-value < 0.001) (Fig. 3d, left panels). Oppositely, Wang and Phillips PNGSs were increased in sg*Fosl1*_1 cells (Wang: NES = 1.42; FDR q-value = 0.029; Phillips: NES = 2.10; FDR q-value < 0.001) (Fig. 3d, right panels). These findings were validated by qRT-PCR with a significant decrease in expression of a panel of MES genes (*Plau*, *Itga7*, *Timp1*, *Plaur*, *Fn1*, *Cyr61*, *Actn1*, *S100a4*, *Vim*, *Cd44*) (Fig. 3e) and increased expression of PN genes (*Olig2*, *Ncam1*, *Bcan*, *Lgr5*) in the *Fosl1* knock-out (KO) *Kras^G12V^* NSCs (Fig. 3f).

### *Fosl1* depletion affects the chromatin accessibility of MES transcription program and differentiation genes

FOSL1 is a member of the AP-1 transcription factor super family, which may be composed of a diverse set of homo- and heterodimers of the individual members of the JUN, FOS, ATF and MAF protein families. In GBM, AP-1 can act as a pioneer factor for other transcriptional regulators, such as ATF3, to coordinate response to stress in GBM ^41^. To test the effect of *Fosl1* ablation on chromatin regulation, we performed open chromatin profiling using ATAC-seq in the p53-null *Kras^G12V^* NSCs model (Fig. 3c). This analysis revealed that *Fosl1* loss strongly affects chromatin accessibility of known cis-regulatory elements such as transcription start sites and CpG islands (TSS and CGI, respectively), as gauged by unsupervised clustering of *Fosl1* wild-type and knock-out cells (Fig. 4a). Consistent with a role for *FOSL1*/FRA-1 in maintaining chromatin accessibility at direct target genes, deletion of *Fosl1* caused the selective closing of chromatin associated with the major AP-1 transcription factors binding sites (Fig. 4b). Profiling of the motifs associated with decreased accessibility upon *Fosl1* loss, returned AP-1/2-mediated functional categories (Fig. 4c). Notably, chromatin that was opened upon *Fosl1* deletion, showed enrichments for a diverse set of general and lineage-specific TFs, including MFZ1, NRF1, RREB1 and others (Fig. 4c). The genes associated with changes in chromatin accessibility upon *Fosl1* loss are involved in several cell fate commitment, differentiation and morphogenesis programs (Fig. 4e,f). Next, we investigated chromatin remodeling dynamics using *limma*, and identified 9749 regions with significant differential accessibility (absolute log2 Fold-change >1, FDR < 0.05). Importantly, *Fosl1* loss induced opening of chromatin associated with lineage-specific markers, along with closing of chromatin at the *loci* of genes, associated with mesenchymal GBM identity in human tumors and BTSC lines (Fig. 4f,g). Taken all together, this evidence further indicates that FRA-1 might modulate the mesenchymal transcriptional program by regulating the chromatin accessibility of MES genes.

**Fig. 4.**
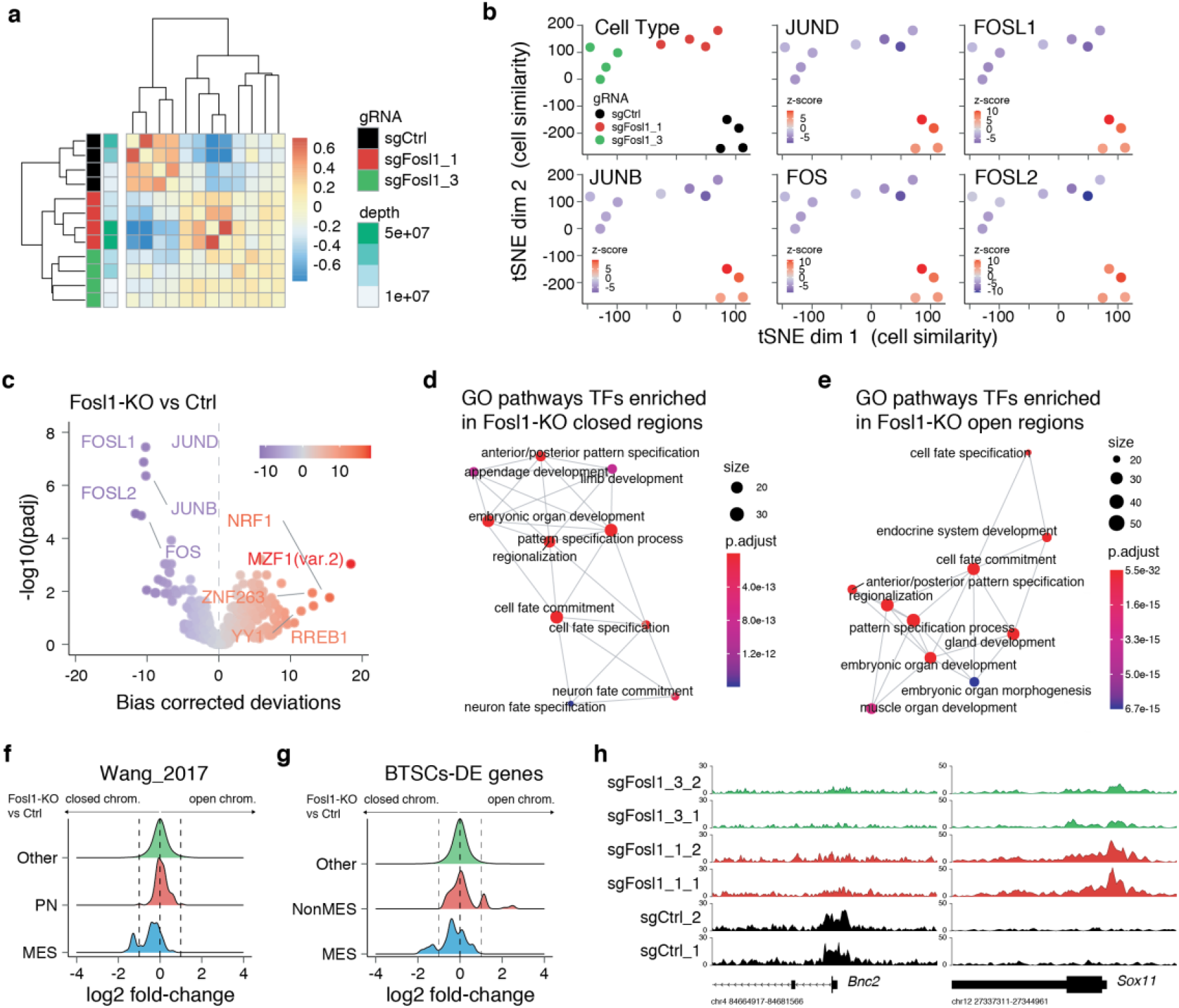
*Fosl1* depletion affects the chromatin accessibility of MES transcription program and differentiation genes in mouse glioma-initiating cells. **a,** Correlation heatmap of the ATAC-seq samples. Clustering of the *Fosl1*-WT (sgCtrl, n = 4) and *Fosl1*-depleted (sg*Fosl1*_1 and sg*Fosl1*_3, n = 8) samples is based upon the bias corrected deviations in chromatin accessibility (see methods). **b,** tSNE visualization of cellular similarity between *Fosl1*-depleted and control cells based on chromatin accessibility. Samples are color-coded according to the cell type (black, red and green for sgCtrl, sgFosl1_1 and sgFosl1_3 cells, respectively), or by directional z-scores. **c,** Volcano plot illustrating the mean difference in bias-corrected accessibility deviations between *Fosl1*-deficient and controls cells against the FDR-corrected p-value for that difference. The top differential motifs are highlighted in violet and red, decreased and increased accessibility respectively. **d**, **e,** Top enriched Gene Ontology (GO) biological processes pathways for the regions with decreased (**d**) and increased (**e**) chromatin accessibility upon *Fosl1* loss. The nodes represent the functional categories from the respective databases, color-coded by the significance of enrichment (FDR <0.05). The node size indicates the number of query genes represented among the ontology term and the edges highlight the relative relationships among these categories. **f, g**, Density plots showing the distributions of the log2 fold-changes in chromatin accessibility of the indicated probes, as measured with *limma* by comparing *Fosl1*-KO versus control cells. **h,** Representative ATAC-seq tracks for the MES *Bnc2* and Non-MES *Sox11* markers loci. Tracks are color-coded as in panel (**a**) and (**b**).

### *Fosl1* deletion reduces stemness and tumor growth

Ras activating mutations have been widely used to study gliomagenesis, in combination with other alterations as *Akt* mutation, loss of *Ink4a/Arf* or *Trp53* ^42–46^. Thus, we then explored the possibility that *Fosl1* could modulate the tumorigenic potential of the p53-null *Kras* mutant cells.

Cell viability was significantly decreased in *Fosl1* knockout cell lines, as compared to sgCtrl (Fig. 5a). Concomitantly, we observed a significant decreased percentage of cells in S-phase (mean values: sgCtrl = 42.6%; sgFosl1_1 = 21.6%, Student’s t test P ≤ 0.001; sgFosl1_3 = 20.4%, Student’s t test P = 0.003), an increase in percentage of cells in G2/M (mean values: sgCtrl = 11.7%, sgFosl1_1 = 28.4%, Student’s t test P ≤ 0.001; sgFosl1_3 = 23.4%, Student’s t test P = 0.012) (Fig. 5b) and a reduction of the expression of cell-cycle regulator genes (*Ccnb1*, *Ccnd1*, *Ccne1* and *Cdk1*, among others) (Extended Data Fig. 5a).

**Fig. 5.**
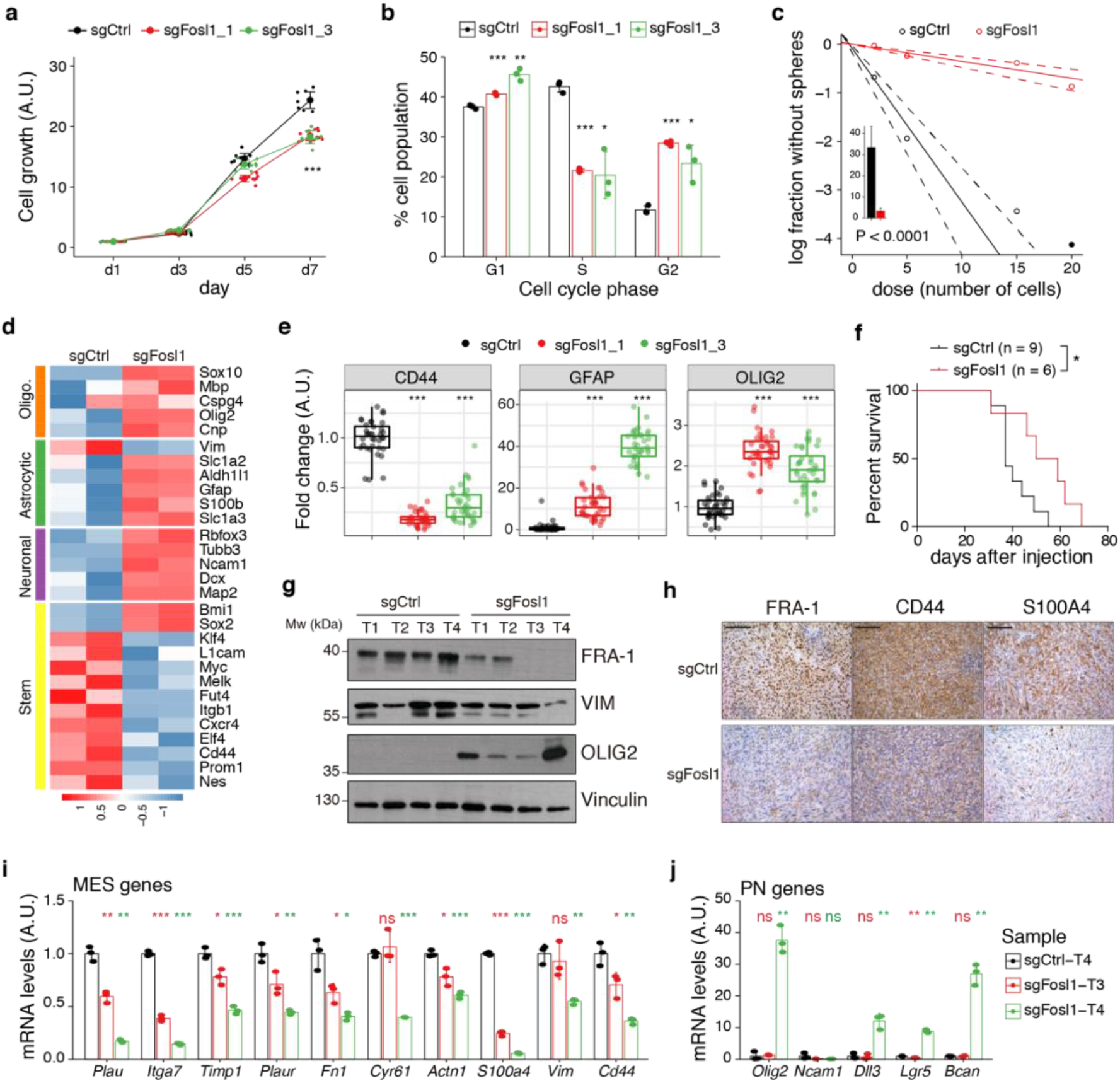
*Fosl1* knock-out impairs cell growth and stemness *in vitro* and increases survival in a orthotopic glioma model. **a,** Cell viability of control and *Fosl1* KO p53-null *Kras^G12V^* NSCs measured by MTT assay; absorbance values were normalized to day 1. Data from a representative of three independent experiments are presented as mean ± SD (n=10). Two-way ANOVA, relative to sgCtrl for both sg*Fosl1*_1 and sg*Fosl1*_3: ***P ≤ 0.001. **b,** Quantification of cell cycle populations of control and *Fosl1* KO p53-null *Kras^G12V^* NSCs by flow cytometry analysis of PI staining. Data from a representative of two independent experiments are presented as mean ± SD (n=3). Student’s t test, relative to sgCtrl: *P ≤ 0.05; **P ≤ 0.01; ***P ≤ 0.001. **c,** Representative limiting dilution experiment on p53-null *Kras^G12V^* sgCtrl and sg*Fosl1*_1 NSCs, calculated with extreme limiting dilution assay (ELDA) analysis; bar plot inlet shows the estimated stem cell frequency with the confidence interval; Chi-square P < 0.0001. **d,** Heatmap of expression of stem cell (yellow) and lineage-specific (neuronal – purple, astrocytic – green and oligodendrocytic – orange) genes, comparing sgCtrl and sg*Fosl1*_1 p53-null *Kras^G12V^* NSCs. **e,** Quantification of pixel area (fold change relative to sgCtrl) of CD44, GFAP and OLIG2 relative to DAPI pixel area per field of view in control and *Fosl1* KO p53-null *Kras^G12V^* NSCs. Data from a representative of two independent experiments; Student’s t test, relative to sgCtrl: ***P ≤ 0.001. **f,** Kaplan-Meier survival curves of *nu*/*nu* mice injected with p53-null *Kras^G12V^* sgCtrl (n=9) and sg*Fosl1*_1 (n=6) NSCs. Log-rank P = 0.0263. **g,** Western blot analysis using the indicated antibodies of 4 sgCtrl and 4 sg*Fosl1*_1 tumors (showing low or no detectable expression of FRA-1); Vinculin used as loading control. **h,** Representative images of IHCs using the indicated antibodies. Scale bars represent 100 μm. **i,** mRNA expression of MES genes in the samples sgCtrl–T4 (higher FRA-1 expression) and sg*Fosl1*_1–T3 and –T4 (no detectable FRA-1 expression). **j,** mRNA expression of PN genes in samples as in (**h**). Data from a representative of two experiments are presented as mean ± SD (n=3), normalized to *Gapdh* expression. Student’s t test for sgFosl1_1 tumors, relative to sgCtrl–T4: ns = not significant, *P ≤ 0.05, **P ≤ 0.01, ***P ≤ 0.001.

Another aspect that contributes to GBM aggressiveness is its heterogeneity, attributable in part to the presence of glioma stem cells. By using limiting dilution assays, we found that *Fosl1* is required for the maintenance of stem cell capacity with a stem cell frequency estimate of sgCtrl = 3 and sgFosl1_1 = 28.6 (chi-square P < 2.2e-16) (Fig. 5c). Moreover, RNA-seq analysis showed that sg*Fosl1*_1 cells downregulated the expression of stem genes (*Elf4*, *Klf4*, *Itgb1*, *Nes*, *Sall4*, *L1cam*, *Melk*, *Cd44*, *Myc*, *Fut4*, *Cxcr4*, *Prom1*) while upregulating the expression of lineage-specific genes: neuronal (*Map2*, *Ncam1*, *Tubb3*, *Slc1a2*, *Rbfox3*, *Dcx*), astrocytic (*Aldh1l1*, *Gfap*, *S100b*, *Slc1a3*) and oligodendrocytic (*Olig2*, *Sox10*, *Cnp*, *Mbp*, *Cspg4*) (Fig. 5d). The different expression of some of the stem/differentiation markers was confirmed also by immunofluorescence analysis. While *Fosl1* KO cells presented low expression of the stem cell marker CD44, differentiation markers as GFAP and OLIG2 were significantly higher when compared to sgCtrl cells (Fig. 5e, Extended Data Fig. 5b).

We then sought to test whether: i) p53-null *Kras^G12V^* NSCs were tumorigenic and ii) *Fosl1* played any role in their tumorigenic potential. Intracranial injections of p53-null *Kras^G12V^* NSCs in *nu*/*nu* mice led to the development of high-grade tumors with a median survival of 37 days in control cells (n=9). However, sg*Fosl1*_1 injected mice (n=6) had a significant increase in median survival (54.5 days, Log-rank *P* = 0.0263) (Fig. 5f). Consistent with what we detected *in vitro* (Fig. 3d-f) we observed a switch from a MGS to a PNGS in the tumors (Fig. 5g-i). By western blot and immunohistochemical analysis, we observed a reduction on expression of MES markers (VIM, CD44 and S100A4) as compared to sgCtrl tumors (Fig. 5g,h), while the PN marker OLIG2 was only found expressed in sg*Fosl1* tumors (Fig. 5g). Similarly, when we compared mRNA expression of a sgCtrl tumor with high FRA-1 expression (T4, Fig. 5g) with sg*Fosl1* tumors with no detectable FRA-1 expression by western blot (T3 and T4, Fig. 5g), we found downregulated expression of MES markers and upregulated expression of PN markers in the sg*Fosl1* tumors (Fig. 5i-j).

Altogether, our data support the conclusion that, besides controlling cell proliferation, *Fosl1* plays a critical role in the maintenance of the stem cell properties and tumorigenicity of p53-null *Kras* mutant NSCs.

### *Fosl1* amplifies MES gene expression

To further assess the role of *Fosl1* as a key player in the control of the MGS, we used a mouse model of inducible *Fosl1* overexpression containing the alleles *Kras^LSLG12V^; Trp53^lox^; ROSA26^LSLrtTA-IRES-EGFP^; Col1a1^TetO-Fosl1^* (here referred as *Fosl1^tetON^*). Similar to the loss-of-function approach here used, this allelic combination allows the expression of *Kras^G12V^* and deletion of *p53* after Cre recombination. Moreover, the expression of the reverse tetracycline transactivator (rtTA) allows, upon induction with doxycycline (Dox), the ectopic expression of *Fosl1* (Flag tagged), under the control of the *Col1a1* locus and a tetracycline-responsive element (TRE or Tet-O)^47,48^.

NSCs derived from *Fosl1^WT^* and *Fosl1^tetON^* mice were infected *in vitro* with a lentiviral vector expressing the Cre recombinase and efficient infection was confirmed by fluorescence microscopy, as the cells expressing the rtTA should express GFP (data not shown). FRA-1 overexpression, as well as Flag-tag expression was then tested by western blot after 72h of Dox induction (Fig. 6a). When *Fosl1^tetON^* NSCs were analyzed by qRT-PCR for the expression of MES/PN markers, a significant upregulation of most MES genes and downregulation of PN genes was found in the cells overexpressing *Fosl1* upon Dox induction (Fig. 6b,c), the inverse image of our findings with *Fosl1* knock-out cells.

**Fig. 6.**
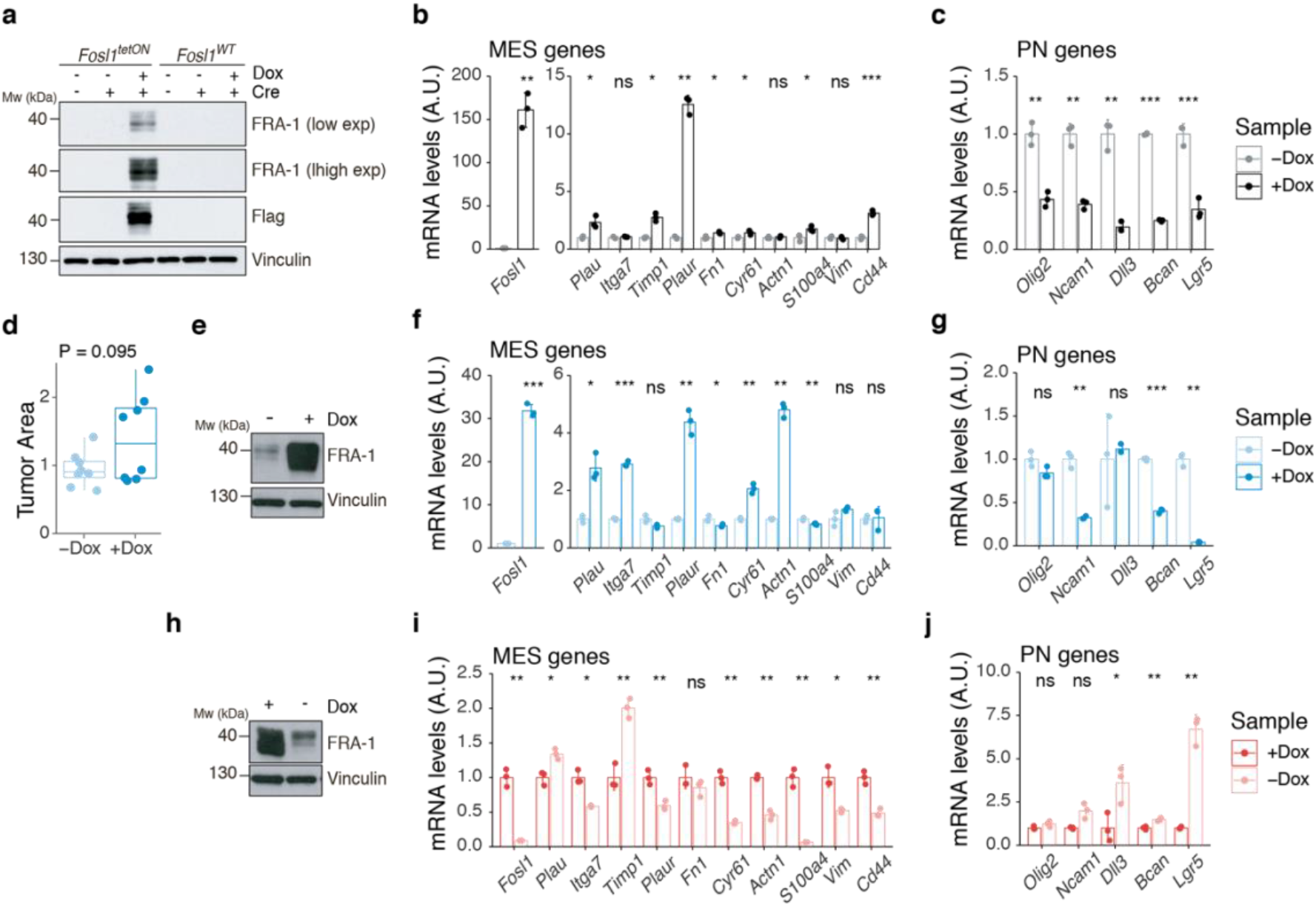
*Fosl1* overexpression upregulates the MGS and induces larger tumors *in vivo*. **a,** Western blot analysis of FRA-1 and Flag expression on *Fosl1^tetON^* and *Fosl1^WT^* NSCs derived from *Kras^LSLG12V^; Trp53^lox^; ROSA26^LSLrtTA-IRES-EGFP^; Col1a1^TetO-Fosl1^* mice, upon *in vitro* infection with Cre and induction of *Fosl1* overexpression with 1 μg/mL Dox for 72 h; Vinculin used as loading control. **b,** mRNA expression of *Fosl1* and MES genes in *Fosl1^tetON^* p53-null *Kras^G12V^* cells upon 72 h induction with 1 μg/mL Dox. **c,** mRNA expression of PN genes in *Fosl1^tetON^* p53-null *Kras^G12V^* cells upon 72 h induction with 1 μg/mL Dox. **d,** Quantification of tumor area (μm^2^) of –Dox and +Dox tumors (n=8/8). For each mouse, the brain section on the H&E slide with a larger tumor was considered and quantified using the ZEN software (Zeiss). **e,** Western blot detection of FRA-1 expression in tumorspheres derived from a control (−Dox) tumor. Tumorspheres were isolated and kept without Dox until first passage, when 1 μg/mL Dox was added and kept for 19 days (+Dox *in vitro*). **f,** mRNA expression of *Fosl1* and MES genes in tumorspheres in absence or presence of Dox for 19 days. **g,** mRNA expression of PN genes in tumorspheres in absence or presence of Dox for 19 days. **h,** Western blot detection of FRA-1 expression in tumorspheres derived from a *Fosl1* overexpressing (+Dox) tumor. Tumorspheres were isolated and kept with 1 μg/mL Dox until first passage, when Dox was removed for 19 days (−Dox *in vitro*). **i,** mRNA expression of *Fosl1* and MES genes in tumorspheres in presence or absence of Dox for 19 days. **j,** mRNA expression of PN genes in tumorspheres in presence or absence of Dox for 19 days. qRT-PCR data from a representative of two experiments are presented as mean ± SD (n=3), normalized to *Gapdh* expression. Student’s t test, relative to the respective control (−Dox in b, c, f and g; +Dox in i and j): ns = not significant, *P ≤ 0.05, **P ≤ 0.01, ***P ≤ 0.001.

To investigate if the MES phenotype induced with *Fosl1* overexpression would have any effect *in vivo*, p53-null *Kras^G12V^ Fosl1^tetON^* NSCs were intracranially injected into syngeneic C57BL/6J wildtype mice. Injected mice were randomized and subjected to Dox diet (food pellets and drinking water) or kept as controls with regular food and drinking water with 1% sucrose. No differences in mice survival were observed (Extended Data Fig. 6a). However, tumors developed from *Fosl1* overexpressing mice (+Dox) were larger (Fig. 6d), more infiltrative and with a more aggressive appearance than controls (–Dox), that mostly grew as superficial tumor masses, even if both –Dox and +Dox tumors seem to proliferate similarly (Extended Data Fig. 6b).

Tumorspheres were derived from –Dox and +Dox tumor-bearing mice and *Fosl1* expression was manipulated *in vitro* through addition or withdrawal of Dox from the culture medium. In the case of tumorspheres derived from a –Dox tumor, when Dox was added for 19 days, high levels of FRA-1 expression were detected by western blot (Fig. 6e). At the mRNA level, Dox treatment also greatly increased *Fosl1* expression, as well as some of the MES genes (Fig. 6f), while the expression of PN genes was downregulated (Fig. 6g). Conversely, when Dox was removed from +Dox derived tumorspheres for 19 days, the expression of FRA-1 decreased (Fig. 6h,i), along with the expression of MES genes (Fig. 6i), while PN genes were upregulated (Fig. 6j). These results confirm the essential role of *Fosl1* in the regulation of the MES gene signature in p53-null *Kras^G12V^* tumor cells and the plasticity between the PN and MES subtypes.

### *FOSL1* controls growth, stemness and MES gene expression in patient-derived tumor cells

To prove the relevance of our findings in the context of human tumors, we analyzed BTSC lines characterized as Non-MES (h676, h543 and BTSC 268) or MES (BTSC 349, BTSC 380 and BTSC 233) (this study and ref ^13^). By western blot, we found that consistent with what observed either in human BTSCs (Fig. 1d) or mouse NSCs (Fig. 3a), MES cell lines expressed high levels of FRA-1 and activation of the MEK/ERK pathway (Fig. 7a).

**Fig. 7.**
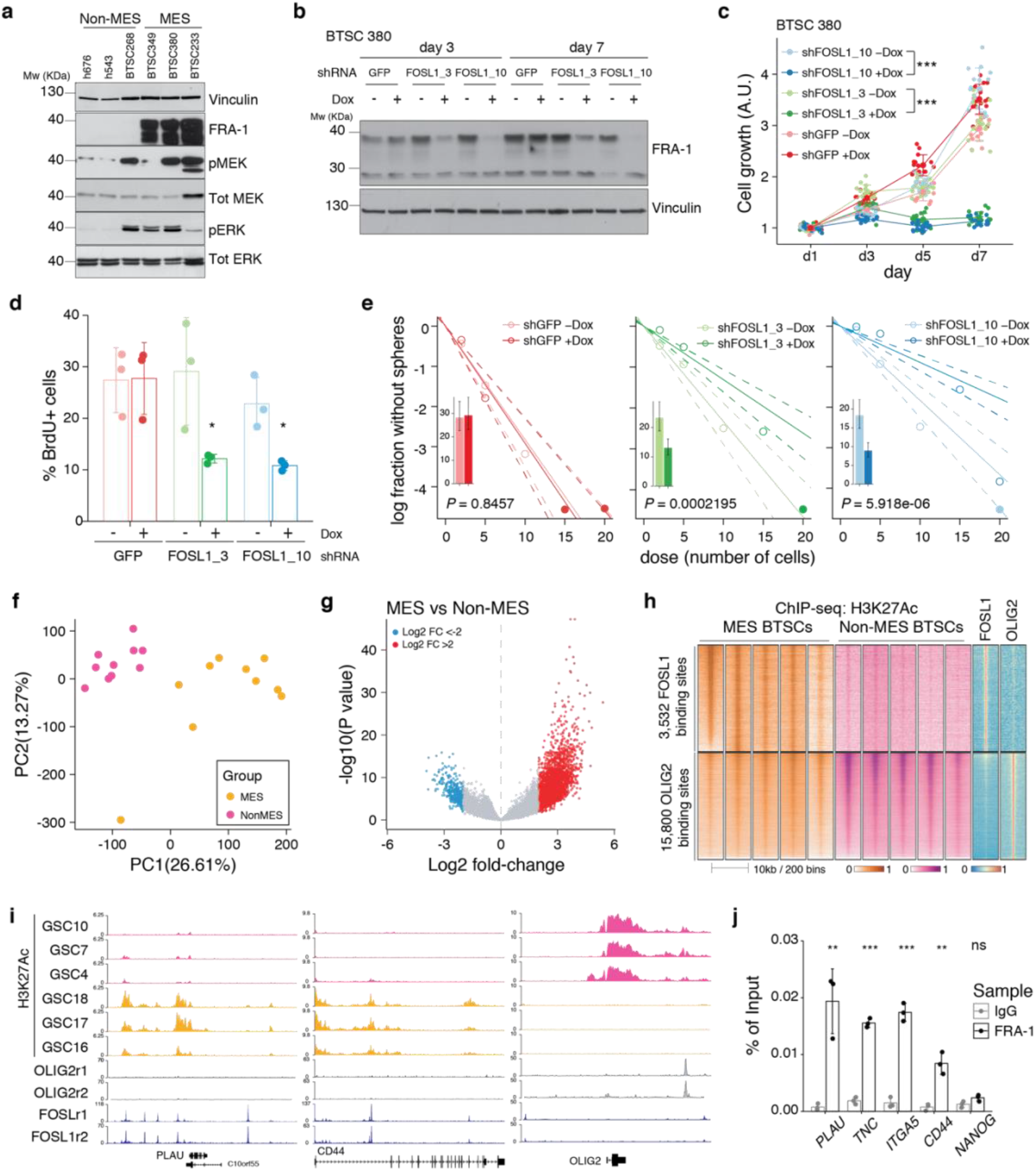
*FOSL1* contributes to MES genes activation, cell growth and stemness in MES BTSCs. **a,** Western blot analysis using the specified antibodies of human brain tumor stem cell lines, characterized as Non-MES (*left*) and MES (*right*). **b,** Western blot detection of FRA-1 in MES BTSC 380 upon transduction with inducible shRNAs targeting GFP (control) and *FOSL1*, analyzed after 3 and 7 days of Dox treatment; Vinculin used as loading control. **c,** Cell growth of BTSC 380 shGFP and sh*FOSL1*, in absence or presence of Dox, measured by MTT assay; absorbance values were normalized to day 1. Data from a representative of three independent experiments are presented as mean ± SD (n=15). Two-way ANOVA, –Dox vs +Dox: ***P ≤ 0.001. **d,** BrdU of BTSC 380 shGFP and sh*FOSL1*, in absence or presence of Dox, analyzed by flow cytometry. Data from a representative of two independent experiments are presented as mean ± SD (n=3). Student’s t test, relative to the respective control (–Dox): *P ≤ 0.05. **e,** Representative limiting dilution analysis on BTSC380 for shGFP and sh*FOSL1*, in presence or absence of Dox, calculated with extreme limiting dilution assay (ELDA) analysis; bar plot inlets show the estimated stem cell frequency with the confidence interval; Chi-square P values are indicated. **f,** Principal component analysis of H3K27Ac signal over *FOSL1*/FRA-1 binding sites, calculated using MACS on ENCODE samples (see methods), in Non-MES (n=10) and MESBTSC (n=10) (from Mack et al., 2019). **g,** Volcano plot illustrating the log2 fold-change differences in H3K27Ac signal between Non-MES and MES BTSCs against the P-value for that difference. Blue and red probes represent statistically significant difference (FDR <0.005) in H3K27Ac signal between Non-Mes and MES BTSCs. **h,** Heatmap of ChIP-seq enrichment of *FOSL1*/FRA-1 or OLIG2 binding sites for the indicated profiles. **i,** View of the *PLAU*, *CD44* and *OLIG2* loci of selected profiles. **j,** Representative ChIP experiment in BTSC 349 cells. The panel shows FRA-1 binding to the promoter of a subset of MES targets (n=3 PCR replicates) expressed as percentage of the initial DNA amount in the immune-precipitated fraction. *NANOG* gene was used as a negative control. Student’s t test, relative to IgG: ns = not-significant, **P ≤ 0.01, ***P ≤ 0.001.

To study the role of *FOSL1* in the context of human BTSCs, its expression was modulated in the MES BTSC 380 using two Dox-inducible shRNAs (sh*FOSL1*_3 and sh*FOSL1*_10). We confirmed by western blot FRA-1 downregulation after 3 and 7 days of Dox treatment (Fig. 7b). In line to what was observed in mouse glioma-initiating cells, *FOSL1* silencing in MES BTSC 380 resulted in reduced cell growth (Fig. 7c) with a significant reduction of the percentage of BrdU positive cells, compared to Dox-untreated cells (Fig. 7d). Moreover, *FOSL1* silencing decreased the sphere forming capacity of MES BTSC 380 with an estimated stem cell frequency of: shGFP –Dox = 3.5, shGFP +Dox = 3.4, chi-square P = 0.8457; sh*FOSL1*_3 –Dox = 4.3, sh*FOSL1*_3 +Dox = 7.6, chi-square P = 0.0002195; sh*FOSL1*_10 –Dox = 5.4, sh*FOSL1*_10 +Dox = 11.1, chi-square P = 5.918e-06 (Fig. 7e). Comparable results were also obtained in the MES BTSC 349 cells (Extended Data Fig. 7a-d). Furthermore, *FOSL1* silencing resulted also in the significant downregulation of the MES genes (Extended Data Fig. 7e, *left panel*), while no major differences in the expression of PN genes was observed (Extended Data Fig. 7e, *right panel*). Lastly, similar to what was observed in mouse tumors (Extended Data Fig. 6b), *FOSL1* overexpression in two Non-MES lines (h543 and h676) did not lead to changes in their proliferation capacity (Extended Data Fig. 7f,g).

We then tested whether FRA-1 modulates the MGS via direct target regulation. To this end, we first identified high-confidence *FOSL1*/FRA-1 binding sites in chromatin immunoprecipitation-seq (ChIP-seq) previously generated in the KRAS mutant HCT116 colorectal cancer cell line (see methods) and then we determined the counts per million reads (CPM) of the enhancer histone mark H3K27Ac in a set of MES (n=10) and non-MES BTSCs (n=10) ^49^, selected based on the highest and lowest *FOSL1* expression, respectively. Principal component analysis (PCA) showed a marked separation of the two groups of BTSCs (Fig. 7f). Differential enrichment analysis by DESeq2 revealed 11748 regions statistically significant (FDR < 0.005) for H3K27Ac at *FOSL1*/FRA-1 binding sites in either MES or non-MES BTSCs (Fig. 7g). Next, we compared H3K27Ac distribution over *FOSL1*/FRA-1 binding sites to that of the Non-MES master regulator *OLIG2* (Fig. 1c). This analysis showed that *FOSL1*/FRA-1 binding sites were systematically decorated with H3K27Ac in MES BTSCs, while the inverse trend was observed at *OLIG2* binding sites (Fig. 7h,i). Validation in an independent MES BTSC line (BTSC 349) by ChIP-qPCR confirmed FRA-1 binding at promoters of some MES genes including *PLAU*, *TNC*, *ITGA5* and *CD44* (Fig. 7j).

Altogether, our data support that *FOSL1*/FRA-1 regulates MES gene expression and aggressiveness in human gliomas via direct transcriptional regulation, downstream of *NF1*.

## Discussion

The most broadly accepted transcriptional classification of GBM was originally based on gene expression profiles of bulk tumors ^3^, which did not discriminate the contribution of tumor cells and TME to the transcriptional signatures. It is now becoming evident that both cell-intrinsic and extrinsic cues can contribute to the specification of the MES subtype ^6,14,50^. Bhat and colleagues had shown that while some of the MES GBMs maintained the mesenchymal characteristics when expanded *in vitro* as BTSCs, some others lost the MGS after few passages while exhibiting a higher PNGS ^14^. These data, together with the evidence that xenografts into immunocompromised mice of BTSCs derived from MES GBMs were also unable to fully restore the MES phenotype, suggested that the presence of an intact TME potentially contributed to the maintenance of a MGS, either by directly influencing a cell-intrinsic MGS or by expression of the TME-specific signature. Recently, the transcriptional GBM subtypes were redefined based on the expression of glioma-intrinsic genes, thus excluding the genes expressed by cells of the TME ^6^. Our master regulator analysis on the BTSCs points to the AP-1 family member *FOSL1* as one of the top transcription factors contributing to the cell-intrinsic MGS. Previous tumor bulk analysis identified a related AP-1 family member *FOSL2*, together with *CEBPB*, *STAT3* and *TAZ*, as important regulators of the MES GBM subtype ^7,8^. While *FOSL1* was also listed as a putative MES master regulator ^8^, its function and mechanism of action have not been further characterized since then. Our experimental data show that *FOSL1* is a key regulator of GBM subtype plasticity and MES transition, and define the molecular mechanism through which *FOSL1* is regulated.

Although consistently defined, GBM subtypes do not represent static entities. The plasticity between subtypes happens at several levels. Besides the referred MES-to-PN change in cultured GSCs compared to the parental tumor ^14^, a PN-to-MES shift often occurs upon treatment and recurrence. Several independent studies comparing matched pairs of primary and recurrent tumors demonstrated a tendency to shift towards a MES phenotype, associated with a worse patient survival, likely as a result of treatment-induced changes in the tumor and/or the microenvironment ^5,6,36^. Moreover, distinct subtypes/cellular states, can coexist within the same tumor ^10,11,50,51^ and targeting these multiple cellular components could result in more effective treatments ^51^.

PN-to-MES transition is often considered an EMT-like phenomenon, associated with tumor progression ^12^. The role of *FOSL1* in EMT has been studied in other tumor types. In breast cancer cells, *FOSL1* expression correlates with mesenchymal features and drives cancer stem cells ^17^ and the regulation of EMT seems to happen through the direct binding of FRA-1 to promoters of EMT genes such as *Tgfb1*, *Zeb1* and *Zeb2* ^28^. In colorectal cancer cells, *FOSL1* was also shown to promote cancer aggressiveness through EMT by direct transcription regulation of EMT-related genes ^29,52^.

It is well established that *NF1* inactivation is a major genetic event associated with the MES subtype ^3,6^. However, this is probably a late event in MES gliomagenesis, as all tumors possibly arise from a PN precursor and just later in disease progression acquire *NF1* alterations that are directly associated with a transition to a MES subtype ^13^. Moreover, *NF1* deficiency has been recently linked to macrophage/microglia infiltration in the MES subtype ^6^. The fact that the enriched macrophage/microglia microenvironment is also able to modulate a MES phenotype suggests that there might be a two-way interaction between tumor cells and TME. The mechanisms of *NF1*-regulated chemotaxis and whether this relationship between the TME and MGS in GBM is causal remain elusive.

Here we provide evidence that manipulation of *NF1* expression levels in patient-derived BTSCs has a direct consequence on the tumor-intrinsic MGS activation and that such activation, can at least in part be mediated by the modulation of *FOSL1*. Among the previously validated MRs, only *CEBPB* appears also to be finely modulated by *NF1* inactivation. This suggests that among the TFs previously characterized (such as *FOSL2*, *STAT3*, *BHLHB2* and *RUNX1*), *FOSL1* and *CEBPB* might play a specific role in the *NF1*-mediated MES transition that occurs in glioma cells with limited or possibly absent effect by the TME. However, whether *FOSL1* contributes also to the putative cross-talk between the TME and the cell-intrinsic MGS, still have to be established.

Furthermore we show that *FOSL1* is a crucial player in glioma pathogenesis, particularly in a MAPK-driven MES GBM context (Fig. 8). Our findings broaden its previously described role in KRAS-driven epithelial tumors, such as lung and pancreatic ductal adenocarcinoma ^24^. *NF1* inactivation results in Ras activation, which stimulates downstream pathways as MAPK and PI3K/Akt /mTOR. RAS/MEK/ERK signaling in turn regulates FRA-1 protein stability ^53,54^. *FOSL1* mRNA expression is then most likely induced by binding of the SRF/Elk1 complex to the serum-responsive element (SRE) on *FOSL1* promoter ^55^. At the same time, FRA-1 can then directly bind to its own promoter to activate its own expression ^29,56^ and those of MES genes. In support of this model (Fig. 8), we observed that both SRF and FOSL1 ChIP-seq peaks on *FOSL1* promoter are decorated with the H3K27Ac active chromatin mark exclusively in MES BTSCs lines (Extended Data Fig. 9). This further generates a feedback loop that induces MGS, increases proliferation and stemness, sustaining tumor growth. FRA-1 requires, for its transcriptional activity, heterodimerization with the AP-1 transcription factors JUN, JUNB or JUND ^57^. Which of the JUN family members participate in the MES gene regulation and whether FRA-1 activates MES gene expression and simultaneously represses PN genes, requires further investigation.

**Fig. 8.**
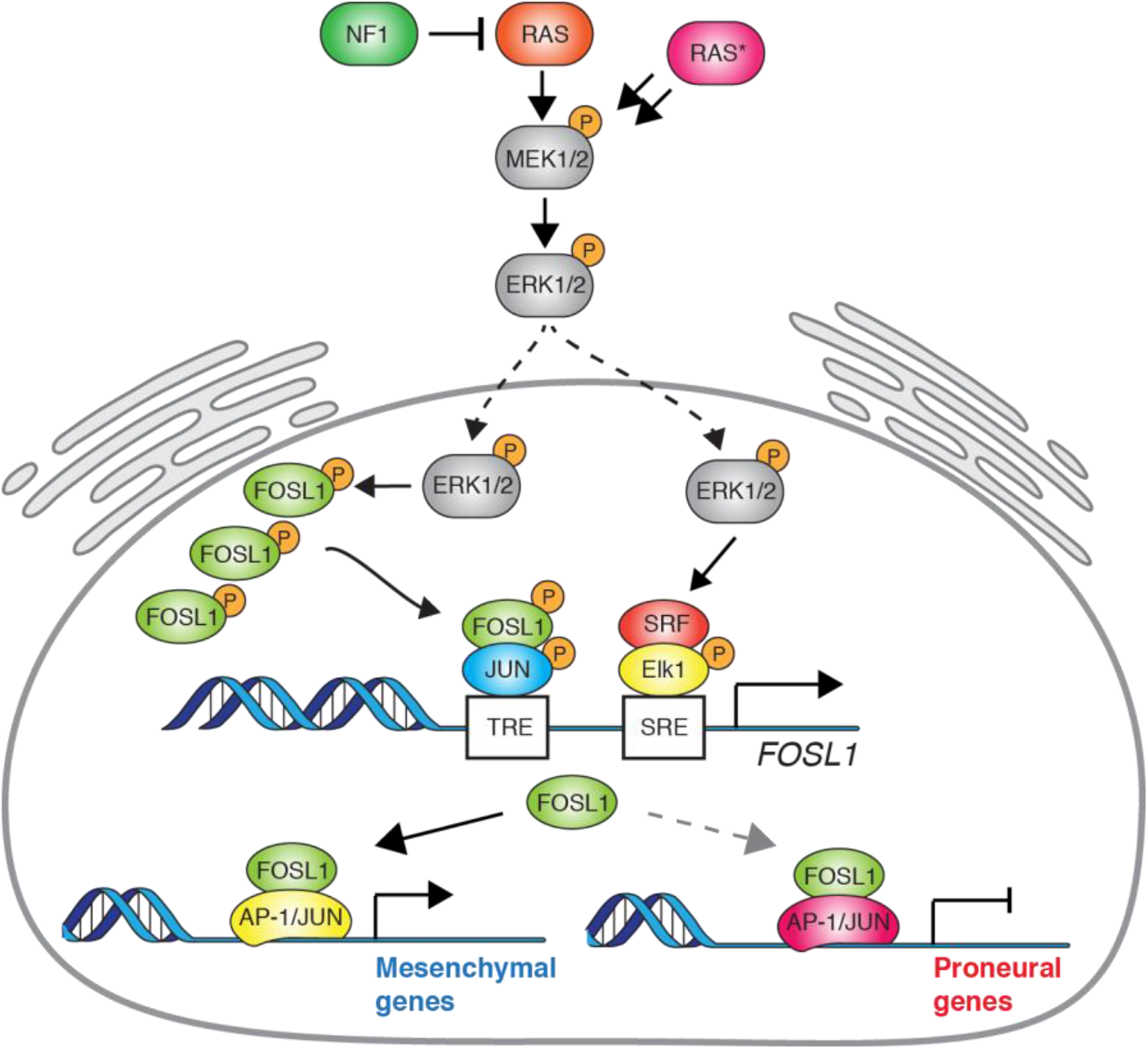
Schematic model of NF1-MAPK-FOSL1 axis in MES gliomas. NF1 alterations or RAS mutations lead to the activation of the MAPK signaling that in turn increases FOSL1 expression both at the mRNA and protein levels. FOSL1 then activates the expression of the MES genes signature and possibly inhibits the PN gene signature. The scheme integrates data presented in this work as well as previously published literature on the regulation of *FOSL1* expression by MAPK activation.

In conclusion, *FOSL1* is a key regulator of the MES subtype of GBM, significantly contributing to its stem cell features, which could open new therapeutic options. Although *FOSL1* pharmacological inhibition is difficult to achieve due to the lack of specific inhibitors, a gene therapy approach targeting *FOSL1* expression through CRISPR, for instance, could constitute an attractive alternative to treat MES GBM patients.

## Acknowledgements

We would like to thank Álvaro Ucero for his input on the project and Flora A. Díaz for her technical support. We are grateful to Francisco X. Real and Scott Lowe for critical input on the manuscript. We thank Pamela Franco for experimental support and discussion. C.M was supported by a “La Caixa” predoctoral fellowship. Y.D. was supported by the Berlin School of Integrative Oncology (BSIO) of the Berlin Charité Medical University. The G.G. lab acknowledges funding from MDC, Helmholtz (VH-NG-1153) and ERC (714922). This work was supported by a grant from the Marie Curie International re-integration Grants (MC-IRG), project nr. 268303 (to M.S.C.) and by grants from the ISCIII, project PI13/01028, cofounded by the European Regional Development Fund (ERDF), and from the Seve Ballesteros Foundation (to M.S.).

## Author Contributions

C.M designed and performed experiments, analyzed data and wrote the manuscript. T.U., P.K., A.I., E.K. and B.O. performed experiments. Y.D. performed and analyzed the ATAC-Seq and contributed to writing the manuscript. G.G. contributed to experimental design, analyzed data, interpreted experiments and edited the manuscript. O.S. provided tumor samples. S.N. provided cell lines. L.B. and E.F.W. provided reagents, contributed to experimental design, interpreted experiments and edited the manuscript. M.S.C and M.S. conceived the project, designed and interpreted experiments and wrote the manuscript.

## Methods

### Generation of the BTSCs dataset and Master regulator analysis (MRA)

The brain tumor stem cell lines (BTSCs) dataset was assembled with new and previously generated transcriptomic data: 22 Illumina HumanHT-12v4 expression BeadChip microarrays newly generated at Freiburg University (GSE137310, this study), 44 RNA-seq samples (Illumina HiSeq 2500) from GSE119834 ^49^, 14 RNA-seq samples (Illumina HiSeq 2000) from SRP057855 ^58^, 30 Affymetrix Human Genome U219 microarrays from GSE67089 ^59^, 17 Affymetrix Human Genome U133 Plus 2.0 microarrays from GSE8049 ^60^, 17 Affymetrix GeneChip Human Genome U133A 2.0 microarrays from GSE49161 ^14^. To analyze the RNA-seq samples we used the Nextpresso pipeline ^61^. For the Affymetrix microarrays, raw data were downloaded from the GEO repository (https://www.ncbi.nlm.nih.gov/geo/) and subsequently the ‘affy’ R package ^62^ was used for robust multi-array average normalization followed by quantile normalization. For genes with several probe sets, the median of all probes had been chosen and only common genes among all the datasets (n = 9889) were used for further analysis. To avoid issues with the use of different transcriptomic platforms each dataset was then scaled (mean = 0, sd = 1) before assembling the combined final dataset. Transcriptional subtypes were obtained using the ‘ssgsea.GBM.classification’ R package ^6^, using 1000 permutations. Differential gene expression (MES vs Non-MES BTSCs) was performed using the ‘limma’ R package ^63^, taking into account the possible batch differences due the different datasets assembled.

The master regulator analysis was performed using the ‘RTN’ R package ^32^. Normalized BTSC expression data were used as input to build a transcriptional network (TN) for 607 TFs present in the dataset. TF annotations were obtained from Gene Ontology (GO:0003700). P values for network edges were computed from a pooled null distribution using 1000 permutations. Edges with an adjusted-P value < 0.05 were kept for data processing inequality (DPI) filtering. In the TN, each target can be connected to multiple TFs and regulation can occur as a result of both direct and indirect interactions. DPI-filtering removes the weakest interaction in any triangle of two TFs and a target gene, therefore preserving the dominant TF-target pairs and resulting in a filtered TN that highlights the most significant interactions ^33^. Post-DPI filtering, the MRA computes the overlap between the transcriptional regulatory unities (regulons) and the input signature genes using the hypergeometric distribution (with multiple hypothesis testing corrections). To identify master regulators, the differential gene expression between MES and Non-MES was used as a phenotype.

### TCGA pan-glioma and CGGA data analysis

RSEM normalized RNA-seq data for the TCGA GBMLGG and CGGA datasets were downloaded from the Broad Institute Firebrowse (http://gdac.broadinstitute.org) and the Chinese Glioma Genome Atlas (updated November 2019) (http://www.cgga.org.cn/), respectively. *NF1* copy number alterations and point mutations for the TCGA GBMLGG were obtained at the cBioPortal (https://www.cbioportal.org). *FOSL1* expression and *NF1* mutational status for the TCGA datasets in Extended Data Fig. 8 were obtained from the Timer2.0 web portal (http://timer.cistrome.org/) ^64^. Transcriptional subtypes were inferred using the ‘ssgsea.GBM.classification’ R package as indicated above. Glioma molecular subtypes information was downloaded from the GlioVis web portal (http://gliovis.bioinfo.cnio.es) ^65^. Survival analysis was performed using the ‘survival’ R package. Stratification of the patients has been done by comparing the 25% FOSL1 high to the rest of the cohort.

### Gene Expression Array and gene set enrichment analysis (GSEA)

For gene expression profiling of the BTSC lines of the Freiburg dataset, total RNA was prepared using the RNeasy kit (Qiagen #74104) or the AllPrep DNA/RNA/Protein mini kit (Qiagen #80004) and quantified using 2100 Bioanalyzer (Agilent). One-and-a-half μg of total RNA for each sample was sent to the genomic facility of the German Cancer Research Center (DKFZ) in Heidelberg (Germany) where hybridization and data normalization were performed. Hybridization was carried out on Illumina HumanHT-12v4 expression BeadChip. Gene set enrichment analysis was performed using the GSEA software (http://www.broadinstitute.org/gsea/index.jsp).

### ATAC-seq

ATAC-seq was performed on 60,000 p53-null Kras^G12V^ mouse neural stem cells transduced with either sgFosl1_1, sgFosl1_3 or sgCtrl. Briefly, the cells were pelleted in PBS and tagmentation was performed in either 50μl of master mix containing 25μL 2×TD buffer, 2.5μL transposase and 22.5μL nuclease-free water (Nextera DNA Library Prep, Illumina #FC-121-1030) or in 50μl of tagmentation mix containing 25μL of TAPS-DMF buffer (80mM TAPS, 40mM MgCl_2_, 50% vol/vol DMF), 0.625μL in-house-produced hyperactive Tn5 enzyme and 24.4μL nuclease-free water (adapted from Hennig et al., 2018). The tagmentation reactions were incubated for 1h at 37°C with moderate agitation (500-800rpm). After the incubation, 5μl of Proteinase K (Invitrogen #AM2548) were added to the samples to stop the transposition. The cells were subsequently lysed by adding 50μl of AL buffer (Qiagen #19075) and incubating for 10 min at 56°C. The DNA was extracted by means of 1.8x vol/vol AMPure XP beads (Beckman Coulter #A63881) and eluted in 18μl. To determine the optimal number of PCR cycles required for library amplification, 2μl of each sample were taken as template for qPCR using KAPA HiFi HotStart ReadyMix (Roche #7958927001) and 1xEvaGreen Dye (Biotium #31000). The whole probe amplification was performed in 50μl qPCR volume with 8-12μl of template DNA. Primers were previously described ^67^. Each library was individually quantified utilizing the Qubit 3.0 Fluorometer (Life Technologies). The appropriate ATAC-seq libraries laddering pattern was determined with TapeStation High Sensitivity D1000 ScreenTapes (Agilent #5067-5584). The libraries were sequenced on the Illumina NextSeq500 using the High Output V2 150 cycles chemistry kit in a 2×75 bp mode.

### ATAC-seq analysis

The reads were adaptor-trimmed using the trim-galore v0.6.2 *--nextera* (https://www.bioinformatics.babraham.ac.uk/projects/trim_galore/). The mapping was conducted using the bowtie2 v2.3.5 ^68^ default parameters. The differential chromatin accessibility analysis was performed in SeqMonk taking mouse CpG islands (mCGI) and Transcription Start Sites (TSSs) ± 500 bp as the probe set (GRCm38 assembly). Counts were normalized by means of the “Read Count Quantification” function with additional correction for total count (CPM), log-transformation and size-factor normalization. The differential accessibility between the *Fosl1*-WT and *Fosl1*-depleted cells was determined utilizing the limma pipeline ^63^.

The “chromVAR” R package ^69^ was used to analyze the chromatin accessibility data, perform corrections for known technical biases and identify motifs with differential deviation in chromatin accessibility between the samples. The enrichGO function from the “clusterProfiler” R package ^70^ was used to visualize the relevant pathways in Fig. 4d and 4e. The bamCoverage function of the deepTools2 tool ^71^ was used to generate BigWig from aligned files for subsequent visualization with the “karyoploteR” R package ^72^.

### ChIP-seq analysis

We downloaded FOSL1 ChIP-seq profiling from the KRAS mutant HCT116 cell line ENCODE tracks ENCFF000OZR and ENCFF000OZQ. FOSL1/FRA-1 peaks (29738) were identified with SeqMonk, using the MACS algorithm ^73^ with a 10^−7^ P value cutoff. OLIG2 binding sites and ChIP-seq profiles were downloaded from GEO: GSM1306365_MGG8TPC.OLIG2r1c and GSM1306367_MGG8TPC.OLIG2r2 ^74^. SRF ChIP-seq profiles were downloaded for the HCT116 ENCODE tracks ENCFF000PCL and ENCFF000PCN. H3K27Ac data were downloaded from GSE119755 (Mack et al., 2019) for: GSM3382275_GSC1_H3K27AC, GSM3382277_GSC10_H3K27AC, GSM3382285_GSC14_H3K27AC, GSM3382289_GSC16_H3K27AC, GSM3382291_GSC17_H3K27AC, GSM3382293_GSC18_H3K27AC, GSM3382295_GSC19_H3K27AC, GSM3382299_GSC20_H3K27AC, GSM3382303_GSC22_H3K27AC, GSM3382313_GSC27_H3K27AC, GSM3382319_GSC3_H3K27AC, GSM3382327_GSC33_H3K27AC, GSM3382331_GSC35_H3K27AC, GSM3382333_GSC36_H3K27AC, GSM3382341_GSC4_H3K27AC, GSM3382343_GSC40_H3K27AC, GSM3382345_GSC41_H3K27AC, GSM3382347_GSC43_H3K27AC, GSM3382351_GSC5_H3K27AC, GSM3382355_GSC7_H3K27AC. Quantitation was Read Count Quantitation using all reads correcting for total count only in probes normalized to the largest store, log transformed and duplicates ignored. Differential H3K27Ac signal was measured through the DEseq2 pipeline ^75^. Heatmaps were generated with ChaSE ^76^, plotting H3K27AC signal over FOSL1 (3532 probes, MES vs Non-MES log2 fold-change >2) or OLIG2 (15800 probes) peaks, with a 10,000 bp probe window.

### Mouse strains and husbandry

*GFAP-tv-a; hGFAP-Cre; Rosa26-LSL-Cas9* mice were previously described (Oldrini et al., 2018)*. Kras^LSLG12V^; Trp53^lox^; Rosa26^LSLrtTA-IRES-EGFP^; Col1a1^TetO-Fosl1^* mouse strain corresponds to the MGI Allele References 3582830, 1931011, 3583817 and 5585716, respectively. Immunodeficient *nu*/*nu* mice (MGI: 1856108) were obtained at the Spanish National Cancer Research Centre Animal Facility.

Mice were housed in the specific pathogen-free animal house of the Spanish National Cancer Research Centre under conditions in accordance with the recommendations of the Federation of European Laboratory Animal Science Associations (FELASA). All animal experiments were approved by the Ethical Committee (CEIyBA) and performed in accordance with the guidelines stated in the International Guiding Principles for Biomedical Research Involving Animals, developed by the Council for International Organizations of Medical Sciences (CIOMS).

### Cell lines, cell culture and drug treatments

Mouse neural stem cells (NSCs) were derived from the whole brain of newborn mice of *Gtv-a; hGFAP-Cre; LSL-Cas9; Trp53^lox^* (referred as p53-null NSCs) and *Kras^LSLG12V^; Trp53^lox^; Rosa26^LSLrtTA-IRES-EGFP^; Col1a1^TetO-Fosl1^* (referred as *Fosl1^TetON^* NSCs). Tumorsphere lines were derived from tumors of C57BL/6J injected with *Fosl1^tetON^* NSCs, when mice were sacrificed after showing symptoms of brain tumor disease. For the derivation of mouse NSCs and tumorspheres, tissue was enzymatically digested with 5 mL of papain digestion solution (0.94 mg/mL papain (Worthington #LS003119), 0.48 mM EDTA, 0.18 mg/mL N-acetyl-L-cysteine (Sigma-Aldrich #A9165) in Earl’s Balanced Salt Solution (Gibco #14155-08)) and incubated at 37°C for 8 min. After digestion, the enzyme was inactivated by the addition of 2 mL of 0.71 mg/mL ovomucoid (Worthington #LS003087) and 0.06 mg/mL DNaseI (Roche #10104159001) diluted in Mouse NeuroCult basal medium (Stem Cell Technologies #05700) without growth factors. Cell suspension was centrifuged at a low speed and then passed through a 40 μm mesh filter to remove undigested tissue, washed first with PBS and then with ACK lysing buffer (Gibco #A1049201) to remove red blood cells. NSCs and tumorspheres were grown in Mouse NeuroCult basal medium, supplemented with Proliferation supplement (Stem Cell Technologies #05701), 20 ng/mL recombinant human EGF (Gibco #PHG0313), 10 ng/mL basic-FGF (Millipore #GF003-AF), 2 μg/mL Heparin (Stem Cell Technologies #07980) and L-glutamine (2mM, Hyclone #SH3003401). Spheres were dissociated with Accumax (ThermoFisher Scientific #00-4666-56) and re-plated every 4-5 days.

Patient-derived glioblastoma stem cells (BTSCs) from Freiburg University were prepared from tumor specimens under IRB-approved guidelines as described before (Fedele et al., 2017). The gender of the main BTSCs lines used in this study are: BTSCT232 (male), BTSCT233 (female), BTSCT349 (female), BTSCT380 (male). BTSCs were grown as neurospheres in Neurobasal medium (Gibco #10888022) containing B27 supplement (Gibco #12587010), N2 supplement (Gibco #17502048), b-FGF (20 ng/mL), EGF (20 ng/mL), LIF (10 ng/mL, CellGS #GFH200-20), 2 μg/mL Heparin and L-glutamine (2mM). Patient-derived glioblastoma stem cell lines h543 and h676, kindly provided by Eric C. Holland, were grown as neurospheres Human NeuroCult basal medium, supplemented with Proliferation supplement (Stem Cell Technologies #05751) and growth factors, as the mouse NSCs or tumorspheres (see above).

The MAPK inhibitors GDC-0623, Trametinib and U0126 were purchased from Merk (AMBH303C5C40), Selleckchem (Cat. S2673) and Sigma-Aldrich (Cat. 662005), respectively.

### Vectors, virus production and infection

Flag-tagged NF1-GRD (aminoacids 1131-1534) was amplified by PCR from human cortical tissue (epilepsy patient) and first cloned in the pDRIVE vector (Qiagen #231124). Primers are listed in Supplementary Table 5. The NF1-GRD sequence was then excised by restriction digestion using PmeI and SpeI enzymes and subcloned in the modified pCHMWS lentiviral vector (kind gift from V. Baekelandt, University of Leuven, Belgium) sites by removing the fLUC region. The correct sequence was verified by sequencing. For *NF1* silencing, sh*NF1* from pLKO (Sigma, TRCN0000238778) (sh*NF1*_1) was subcloned in pGIPZ lentiviral vector (Open Biosystems). The corresponding short hairpin sequence was synthetized (GATC) and amplified by PCR using XhoI and EcoRI sites containing primers. The PCR product was digested using XhoI and EcoRI and subcloned into the pGIPZ vector previously digested with XhoI and PmeI following by digestion with EcoRI. The two vector fragments were ligated with *NF1* short hairpin fragment. The correct insertion and sequence was validated by sequencing. In addition, experiments were performed using shNF1-pGIPZ clone V2LHS_76027 (sh*NF1*_4) and V2LHS_260806 (sh*NF1*_5).

RCAS viruses (RCAS-shNf1, RCAS-sgNf1 and RCAS-Kras^G12V^) used for infection of p53-null NSCs were obtained from previously transfected DF1 chicken fibroblasts (ATCC #CRL-12203) using FuGENE 6 Transfection reagent (Promega #E2691), according to manufacturer’s protocol. DF1 cells were grown at 39°C in DMEM containing GlutaMAX™ (Gibco #31966-021) and 10% FBS (Sigma-Aldrich #F7524).

The pKLV-U6gRNA-PGKpuro2ABFP was a gift from Dr. Kosuke Yusa (Wellcome Sanger Institute) (Addgene plasmid #50946). For cloning of single gRNAs, oligonucleotides containing the BbsI site and the specific gRNA sequences were annealed, phosphorylated and ligated into the pKLV-U6gRNA(BbsI)-PGKpuro2ABFP previously digested with BbsI. Single gRNAs to target *Fosl1* were designed with Guide Scan (http://www.guidescan.com/) and the sequences cloned were sg*Fosl1*_1: TACCGAGACTACGGGGAACC; sg*Fosl1*_2: CCTAGGGCTCGTATGACTCC; sg*Fosl1*_3: ACCGTACGGGCTGCCAGCCC. These vectors and a non-targeting sgRNA control were used to transduce p53-null *Kras^G12V^* NSCs.

The pLVX-Cre and respective control vector were kindly provided by Dr. Maria Blasco (CNIO) and used to transduce *Fosl1^tetON^* NSCs; pSIN-EF1-puro-FLAG-FOSL1 pLKO.1-TET-sh*FOSL1*_3 and sh*FOSL1*_10 and respective control vectors were a gift from Dr. Silve Vicent (CIMA, Navarra University); pBabe-FOSL1 was previously described ^77^.

Gp2-293 packaging cell line (Clontech #631458) was grown in DMEM (Sigma-Aldrich #D5796) with 10% FBS. Lentiviruses generated in this cell line were produced using calcium-phosphate precipitate transfection and co-transfected with second-generation packaging vectors (pMD2G and psPAX2). High-titer virus was collected at 36 and 60 h following transfection.

All cells were infected with lenti- or retroviruses by four cycles of spin infection (200 × g for 2 h), in presence of 8 μg/mL polybrene (Sigma-Aldrich #H9268). Transduced cells were selected after 48 h from the last infection with 1 μg/mL Puromycin (Sigma-Aldrich #P8833).

### Generation of murine gliomas

p53-null *Kras^G12V^* NSCs (5×10^5^ cells) were injected intracranially into 4 to 5 weeks-old immunodeficient *nu*/*nu* male and female mice.

*Fosl1^TetON^* NSCs (5×10^5^ cells) were intracranially injected into 4 to 5 weeks-old wildtype C57Bl/6J female mice that were fed *ad libitum* with 2 g/kg doxycycline-containing pellets. Due to the limited penetration of the blood brain barrier and to insure enough Dox was reaching the brain, 2 mg/mL Dox (PanReac AppliChem #A29510025) was also added to drinking water with 1% sucrose (Sigma-Aldrich #S0389) ^78,79^. Control mice were kept with regular food and 1% sucrose drinking water.

Mice were anaesthetized with 4% isofluorane and then injected with a stereotactic apparatus (Stoelting) as previously described ^80^. After intracranial injection, all mice were routinely checked and sacrificed when developed symptoms of disease (lethargy, poor grooming, weight loss and macrocephaly).

### Immunohistochemistry

Tissue samples were fixed in 10% formalin, paraffin-embedded and cut in 3 μm sections, which were mounted in Superfrost Plus microscope slides (Thermo Scientific #J1810AMNZ) and dried. Tissues were deparaffinized in xylene and re-hydrated through graded concentrations of ethanol in water, ending in a final rinse in water.

For histopathological analysis, sections were stained with hematoxylin and eosin (H&E).

For immunohistochemistry, deparaffinized sections underwent heat-induced antigen retrieval, endogenous peroxidase activity was blocked with 3% hydrogen peroxide (Sigma-Aldrich #H1009) for 15 min and slides were then incubated in blocking solution (2.5% BSA (Sigma-Aldrich #A7906) and 10% Goat serum (Sigma-Aldrich #G9023), diluted in PBS) for at least 1 hr. Incubations with anti-FRA-1 (Santa Cruz #sc-183, 1:100) and anti-CD44 (BD Biosciences #550538, 1:100) were carried out overnight at 4°C. Slides were then incubated with secondary anti-rabbit (Vector #BA-1000) or anti-rat (Vector #BA-9400) for 1 hr at RT and with AB (avidin and biotinylated peroxidase) solution (Vectastain Elite ABC HRP Kit, Vector, PK-6100) for 30 min. Slides were developed by incubation with peroxidase substrate DAB (Vector SK-4100) until desired stain intensity. Finally, slides were counterstained with hematoxylin, cleared and mounted with a permanent mounting medium.

Immunohistochemistry for S100A4 (Abcam #ab27957, 1:300) and Ki67 (Master Diagnostica #0003110QD, undiluted) was performed using an automated immunostaining platform (Ventana discovery XT, Roche).

### Immunoblotting

Cell pellets or frozen tumor tissues were lysed with JS lysis buffer (50 mM HEPES, 150 mM NaCl, 1% Glycerol, 1% Triton X-100, 1.5 mM MgCl2, 5 mM EGTA) and protein concentrations were determined by DC protein assay kit II (Bio-Rad #5000112). Proteins were separated on house-made SDS-PAGE gels and transferred to nitrocellulose membranes (Amersham #10600003). Membranes were incubated in blocking buffer (5% milk in TBST) and then with primary antibody overnight at 4°C. The following primary antibodies and respective dilutions were used: FLAG (Cell Signaling Technology #2368S, 1:2000), FRA-1 (Santa Cruz #sc-183, 1:1000; #sc-605, 1:1000), GFAP (Sigma-Aldrich #G3893, 1:5000), NF1 (Santa Cruz #sc-67, 1:500; Bethyl #A300-140A, 1:1000), OLIG2 (Millipore #AB9610, 1:2000), VIMENTIN (Cell Signaling Technology #5741, 1:3000), p-ERK1/2 (T202/Y204) (Cell Signaling Technology, #9101, 1:2000/3000; Assay Designs #ADI-905-651, 1:250), ERK1/2 (Cell Signaling Technology, #9102, 1:1000; Abcam #ab17942, 1:1000), p-MEK (S217/221) (Cell Signaling Technology, #9154, 1:500/1000), MEK (Cell Signaling Technology, #9122 1:1000), CHI3L1 (Qidel #4815, 1:1000), p85 (Millipore #06-195, 1:10000), VINCULIN (Sigma-Aldrich #V9131, 1:10000) and α-TUBULIN (Abcam #ab7291, 1:10000). Anti-mouse or rabbit-HRP-conjugated antibodies (Jackson ImmunoResearch, #115-035-003 and #111-035-003) were used to detect desired protein by chemiluminescence with ECL Detection Reagent (Amersham, #RPN2106).

### Reverse transcription quantitative PCR

RNA from NSCs and frozen tissue was isolated with TRIzol reagent (Invitrogen #15596-026) according to the manufacturer’s instructions. For reverse transcription PCR (RT-PCR), 500 ng of total RNA was reverse transcribed using the High Capacity cDNA Reverse Transcription Kit (Applied Biosystems #4368814). Quantitative PCR was performed using the SYBR Select Master Mix (Applied Biosystems #4472908) according to the manufacturer’s instructions. qPCRs were run and the melting curves of the amplified products were used to determine the specificity of the amplification. The threshold cycle number for the genes analyzed was normalized to GAPDH. Mouse and human primer sequences are listed in Supplementary Table 5.

RNA from BTSC cells was prepared using the RNeasy kit or the AllPrep DNA/RNA Protein Mini Kit and used for first strand cDNA synthesis using random primers and SuperscriptIII reverse transcriptase (Life Technologies #18080-085). Primer sequences used in qRT-PCR with SYBR Green are listed in Supplementary Table 5. Quantitative real-time PCR (qRT-PCR) STAT3 and CEBPB were performed using pre-validated TaqMan assays (Applied Biosystems): STAT3: Hs01047580, CEBPB: Hs00270923 and 18s rRNA: Hs99999901.

### MTT assay

Cells were seeded in 96-well culture plates (1000 cells per well, 10 wells per cell line) and grown for 7 days. At each timepoint (days 1, 3, 5 and 7), cell viability was determined by MTT assay. Briefly, 10 μL of 5 mg/mL MTT (Sigma-Aldrich #M5655) was added to each well and cells were incubated for 4 h before lysing with a formazan solubilization solution (10% SDS in 0.01 M HCl). Colorimetric intensity was quantified using a plate reader at 590 nm. Values were obtained after subtraction of matched blanks (medium only).

### Cell cycle analysis: Propidium iodide (PI) staining

Cells were harvested and washed twice with PBS prior to fixation with 70% cold ethanol, added drop-wise to the cell pellet while vortexing. Fixed cells were then washed, first with 1% BSA in PBS, then with PBS only and stained overnight with 50 μg/mL PI (Sigma-Aldrich #P4170) and 100 μg/mL RNase A (Roche #10109142001) in PBS. Samples were acquired in a FACSCanto II cytometer (BD Biosciences) and data were analyzed using FlowJo software.

### BrdU incorporation

Cells were pulse-labelled with 10 μM BrdU (Sigma-Aldrich #B9285) for 2 h, harvested and washed twice with PBS prior to fixation with 70% ethanol cold ethanol, added drop-wise to the cell pellet while vortexing. DNA denaturation was performed by incubating samples for 10 min on ice with 0.1 M HCl with 0.5% Tween-20 and samples were then resuspended in water and boiled at 100°C for 10 min. Anti-BrdU-FITC antibody (BD Biosciences #556028) was incubated according to manufacturer’s protocol. After washing with PBSTB (PBS with 0.5% Tween-20 and 1% BSA), samples were resuspended in 25 μg/mL PI and 100 μg/mL RNase A diluted in PBS. Samples were acquired in a FACSCanto II cytometer (BD Biosciences) and data were analyzed using FlowJo software.

### Immunofluorescence

Cells were plated in laminin-coated coverslips and fixed with 4% PFA for 15 min. Cells were then permeabilized with 0.1% Triton X-100 in 0.2% BSA and coverslips were washed and blocked with 10% donkey serum in 0.2% BSA for 1 h. The following primary antibodies were incubated overnight at 4°C: CD44 (BD Biosciences #550538, 1:100), GFAP (Millipore #MAB360, 1:400) and OLIG2 (Millipore #AB9610, 1:100). Secondary antibodies at 1:400 dilution (from Invitrogen, Alexa-Fluor anti-rabbit-488, anti-mouse-488 and anti-rat 594) were incubated for 1 h at RT and after washing coverslips were incubated for 4 min with DAPI (1:4000, Sigma-Aldrich #D8417) and mounted with ProLong Gold Antifade reagent (Invitrogen #P10144).

Fluorescence signal was quantified as the ratio of green/red pixel area relative to DAPI pixel area per field of view, in a total of 36 fields per condition analyzed.

### Neurosphere formation assay and limiting dilution analysis

Neurospheres were dissociated and passed through a 40 μm mesh filter to eliminate non-single cells. Decreasing cell densities were plated in ultra-low attachment 96-well plates (Corning #CLS3474) and fresh medium was added every 3-4 days. The number of positive wells for presence of spheres was counted 2 weeks after plating. Limiting dilution analysis was performed using ELDA R package (http://bioinf.wehi.edu.au/software/elda/).

### RNA-sequencing and analysis on mouse NSCs

One microgram of total RNA from the samples was used. cDNA libraries were prepared using the “QuantSeq 3‘ mRNA-Seq Library Prep Kit (FWD) for Illumina” (Lexogen #015) by following manufacturer instructions. Library generation is initiated by reverse transcription with oligo(dT) priming, and a second strand synthesis is performed from random primers by a DNA polymerase. Primers from both steps contain Illumina-compatible sequences. Adapter-ligated libraries were completed by PCR, applied to an Illumina flow cell for cluster generation and sequenced on an Illumina HiSeq 2500 by following manufacturer’s protocols. Sequencing read alignment and quantification and differential gene expression analysis was performed in the Bluebee Genomics Platform, a cloud-based service provider (www.bluebee.com). Briefly, reads were first trimmed with bbduk from BBTools (BBMap – Bushnell B, https://sourceforge.net/projects/bbmap/) to remove adapter sequences and polyA tails. Trimmed reads were aligned to the GRCm38/mm10 genome assembly with STAR v 2.5 ^81^. Read counting was performed with HTSeq ^82^. Differential gene expression analysis was performed with DESeq2 ^75^. The list of stem/differentiation markers was compiled by combining a previously described gene list ^83^ with other markers ^84^. GSEAPreranked ^85^ was used to perform gene set enrichment analysis of the described indicated signatures on a pre-ranked gene list, setting 1000 gene set permutations.

### Osteogenesis Differentiation Assay

The osteogenesis differentiation assay was performed using the StemPro Osteogenesis Differentiation Kit (Life Technologies #A1007201) according to the manufacturer’s instructions. Briefly, 5×10^3^ cells/cm^2^ were seeded on laminin-coated glass coverslips in a 24-well cell culture plate. Cells were incubated in MSC Growth Medium at 37°C, 5% CO_2_ for 21 days, replacing the medium every 4 days. Cells were then fixed with 4% formaldehyde, stained with Alizarin Red S solution (pH 4.2) and mounted on microscope slides. Pictures were acquired using an Axiovert Microscope (Zeiss).

### Active Ras pull down assay

Active Ras pull down assay was performed using Active Ras pull down assay kit (ThermoFisher Scientific #16117) according to the manufacturer’s instructions. Briefly, cells were grown in 10 cm plates at 80-90% confluency in presence or absence of growth factors (EGF, FGF and LIF), and lysed with the provided buffer. Lysates were incubated with either GDP or GTP for 30 min followed by precipitation with GST-Raf1-RBD. Western blot was performed with the provided anti-RAS antibody (1:250).

### Chromatin preparation and FRA-1 ChIP

BTSC cells were collected at 2×10^6^ cells confluency, washed in PBS and fixed by addition of 1% formaldehyde for 20 min at room temperature. The cells were resuspended in 2 mL Lysis Buffer (50 mM Tris pH 7.5; 1 mM EDTA pH 8.0; 1% Triton; 0.1% Na-deoxycholate; 150 mM NaCl; protease inhibitors) on ice for 20 min. The suspension was sonicated in a cooled Bioruptor Pico (Diagenode), and cleared by centrifugation for 10 min at 13000 rpm. The chromatin (DNA) concentration was quantified using NanoDrop (Thermo Scientific) and the sonication efficiency monitored on an agarose gel. Protein A/G plus-agarose beads (Santa Cruz #sc-2003) were blocked with sonicated salmon sperm (ThermoFisher #AM9680, 200 mg/mL beads) and BSA (NEB #B9000S, 250 mg/mL beads) in dilution buffer (0.5% NP40; 200 mM NaCl; 50 mM Tris pH 8.0; protease inhibitors) for 2 h at room temperature. The chromatin was pre-cleared with 80 μL of blocked beads for 1 hr at 4°C. Pre-cleared chromatin was incubated with 5 μg of FRA-1 antibody (Santa Cruz #sc-605) overnight at 4°C, then with 40 μL of blocked beads for further 2 hr at 4°C. Control mock immunoprecipitation was performed with blocked beads. The beads were washed 1× with Wash1 (20 mM Tris pH 7.5; 2 mM EDTA pH 8.0; 1% Triton; 0.1% SDS; 150 mM NaCl), 1× with Wash2 (20 mM Tris pH 7.5; 2 mM EDTA pH 8.0; 1% Triton; 0.1% SDS; 500 mM NaCl), 1× with LiCl Wash (20 mM Tris pH 7.5; 1 mM EDTA pH 8.0; 1% NP40; 1% deoxycholate; 250 mM LiCl) and 2× in TE (10 mM Tris pH 7.5; 1 mM EDTA). The immuno-complexes were eluted by two 15 min incubations at 30°C with 100 μL Elution buffer (1% SDS, 100 mM NaHCO_3_), and de-crosslinked overnight at 65°C in the presence of 10 U RNase A (Ambion #AM9780). The immune-precipitated DNA was then purified with the QIAquick PCR purification kit (Qiagen #28104) according to manufacturer’s protocol and analyzed by quantitative real-time PCR.

### Statistical analysis

All statistical analyses were performed using R programming language (3.6.3). Statistical differences between groups were assessed by one-way ANOVA, two-way ANOVA or unpaired two-tailed Student’s t tests, unless otherwise specified.

Kaplan–Meier survival curves were produced with GraphPad Prism and P values were generated using the Log-Rank statistics.

Results are presented as mean ± standard deviation (SD), and statistical significance was defined as P ≤ 0.05 for a 95% confidence interval.

### Data and code availability

The accession numbers for data reported in this paper are GEO: GSE137310 (Freiburg BTSCs) and GSE138010 (mouse NSCs). All the R code used for data analysis and plots generation is available at: https://github.com/squatrim/Marques2019.

---

**Extended Data Fig. 1.**
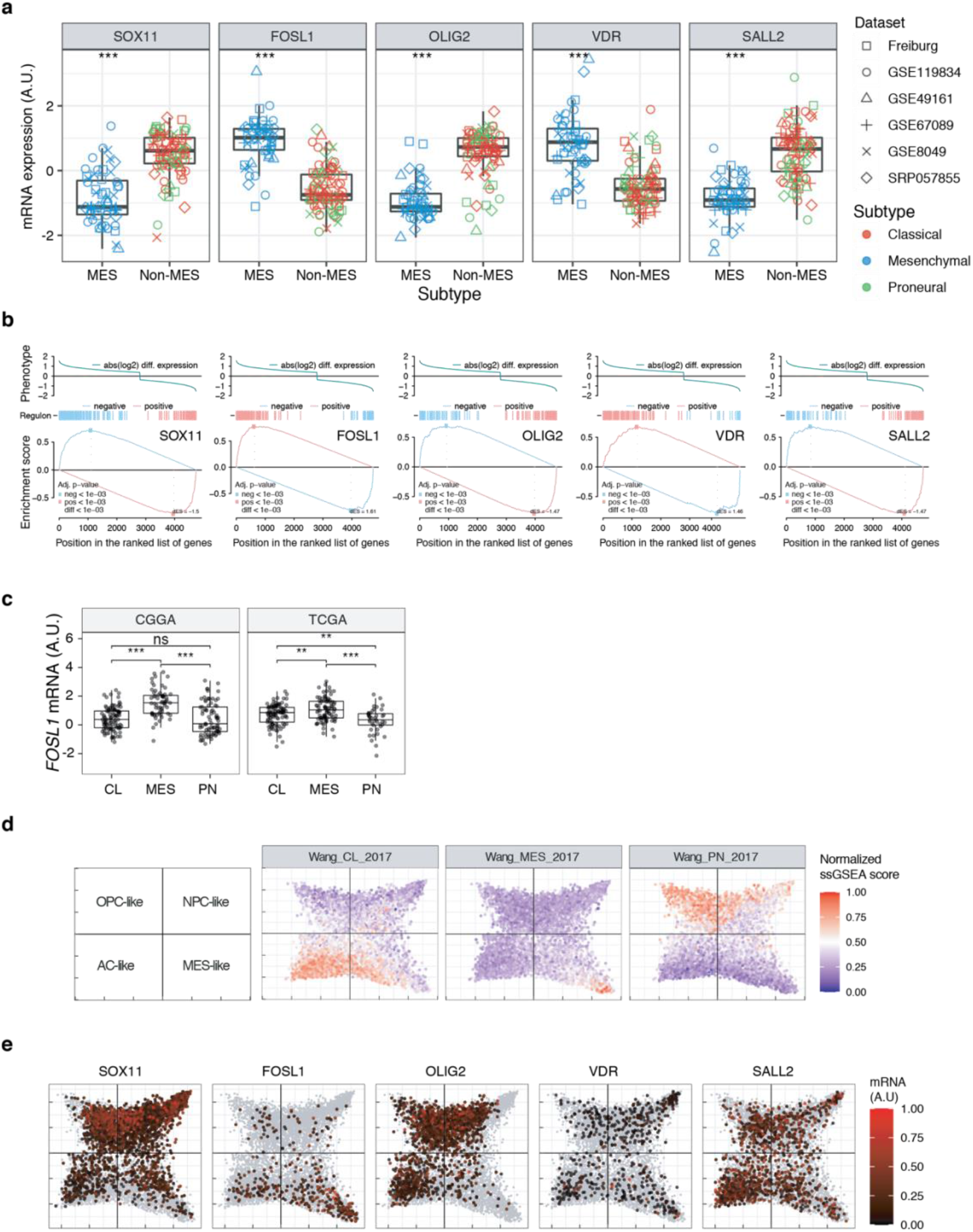
Expression of the top 5 TFs identified in the MRA. **a,** mRNA expression of the top 5 scoring TFs in the MRA of the BTSCs dataset, comparing MES versus Non-MES. Student’s t test, ***P ≤ 0.001. **b,** Two-tailed GSEA showing positive or negative targets for the top 5 TFs in the MRA ranked by their differential expression (MES vs Non-MES). **c,** *FOSL1* mRNA expression in IDH-wt tumors of the CGGA and TCGA datasets. Tumors were separated according to their expression subtype classification. One-way ANOVA with Tukey multiple pairwise-comparison, ***P ≤ 0.001, **P ≤ 0.01, ns = not significant. **d,** ssGSEA scores of the Wang_2017 transcriptional GBM subtypes performed on the scRNA-seq data from Neftel et al 2019. *Left panel*, four main cellular states represented by the four quadrants. **e,** mRNA expression of the top 5 scoring TFs on the scRNA-seq data from Neftel et al. 2019.

**Extended Data Fig. 2.**
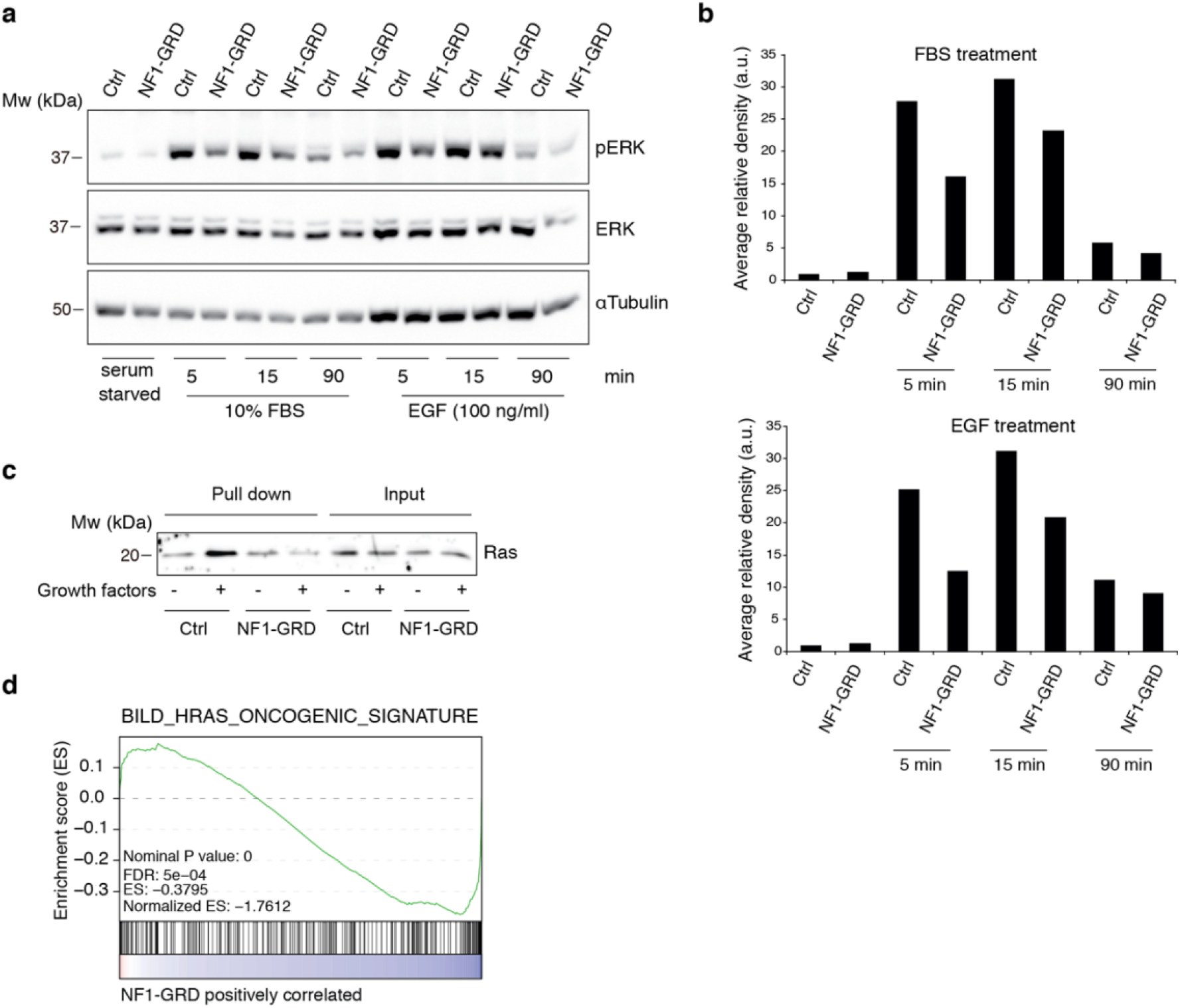
NF1-GRD expression leads to downregulation of RAS signaling. **a,** Western blot analysis of ERK and pERK expression in BTSC 233 cells transduced with NF1-GRD expressing lentivirus and stimulated with 10% FBS or 100 ng/ml EGF. α-Tubulin is included as loading control. **b,** Densitometric analysis of western blot in (**a**). **c,** Western blot analysis of active Ras pull down assay in BTSC 233 expressing NF1-GRD or control, in presence or absence of growth factors. **d,** GSEA of Ras-induced oncogenic signature in BTSC 233 cells transduced with NF1-GRD expressing lentivirus versus Ctrl.

**Extended Data Fig. 3.**
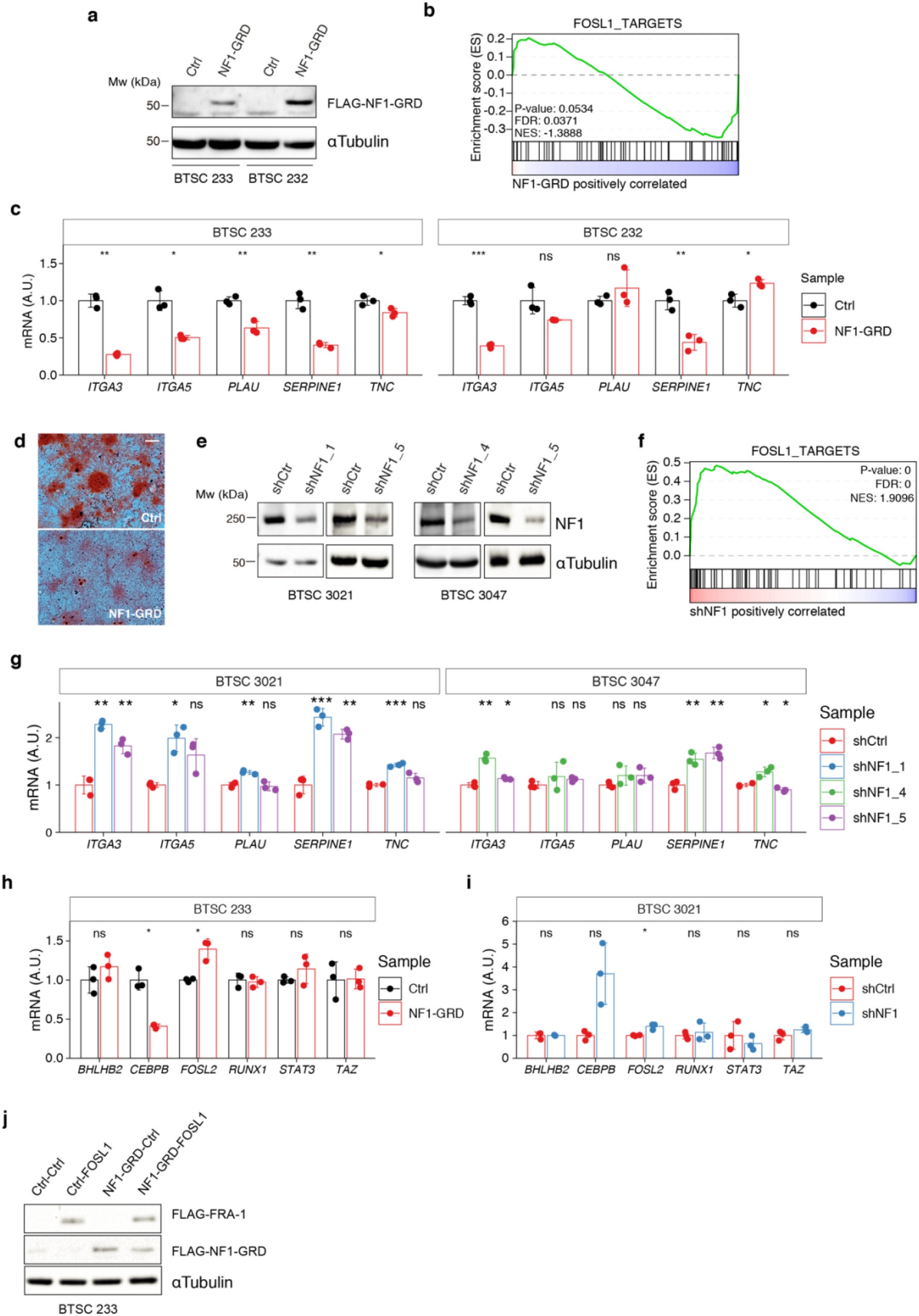
Modulation of NF1 expression regulates FOSL1 targets and mesenchymal genes. **a,** Western blot analysis of FLAG-NF1-GRD expression in MES cells (BTSC 233 and 232). **B,** GSEA of *FOSL1* targets signature in BTSC 233 cells transduced with NF1-GRD or Ctrl vector. **c,** qRT-PCR analysis of mesenchymal *FOSL1* targets (*ITGA3*, *ITGA5*, *PLAU*, *SERPINE1* and *TNC*) in BTSC 233 and 232 cells transduced with NF1-GRD expressing lentivirus. Data are normalized to 18s rRNA expression. **d,** Osteogenesis differentiation assay of BTSC 233 transduced as indicated above. Alzarin Red staining indicates osteogenesis differentiation. Scale bar represents 200 μm. **e,** Western blot analysis of NF1 expression upon *NF1* knockdown in Non-MES cells (BTSC 3021 and 3047). **f,** GSEA of *FOSL1* targets signature in BTSC 3021 cells transduced with sh*NF1* or shCtrl. **g,** qRT-PCR analysis of mesenchymal *FOSL1* targets BTSC 3021 and 3047 cells transduced with sh*NF1* expressing lentiviruses. Data are normalized to 18s rRNA expression. **h, i,** qRT-PCR analysis of MES genes master regulators expression (*BHLHB2*, *CEBPB*, *FOSL2*, *RUNX1*, *STAT3* and *TAZ*) upon NF1-GRD overexpression in BTSC 233 (**h**) or *NF1* knockdown in BTSC 3021 cells (**i**). Data are normalized to GAPDH or 18s rRNA expression, respectively. **j)** Western blot analysis of FLAG-NF1-GRD and FLAG-FRA-1 expression in BTSC 233 cells. qRT-PCR data in (**c**), (**g**), (**f**) and (**h**) are presented as mean ± SD (n=3); Student’s t test, ns = not-significant, *P ≤ 0.05, **P ≤ 0.01, ***P ≤ 0.001.

**Extended Data Fig. 4.**
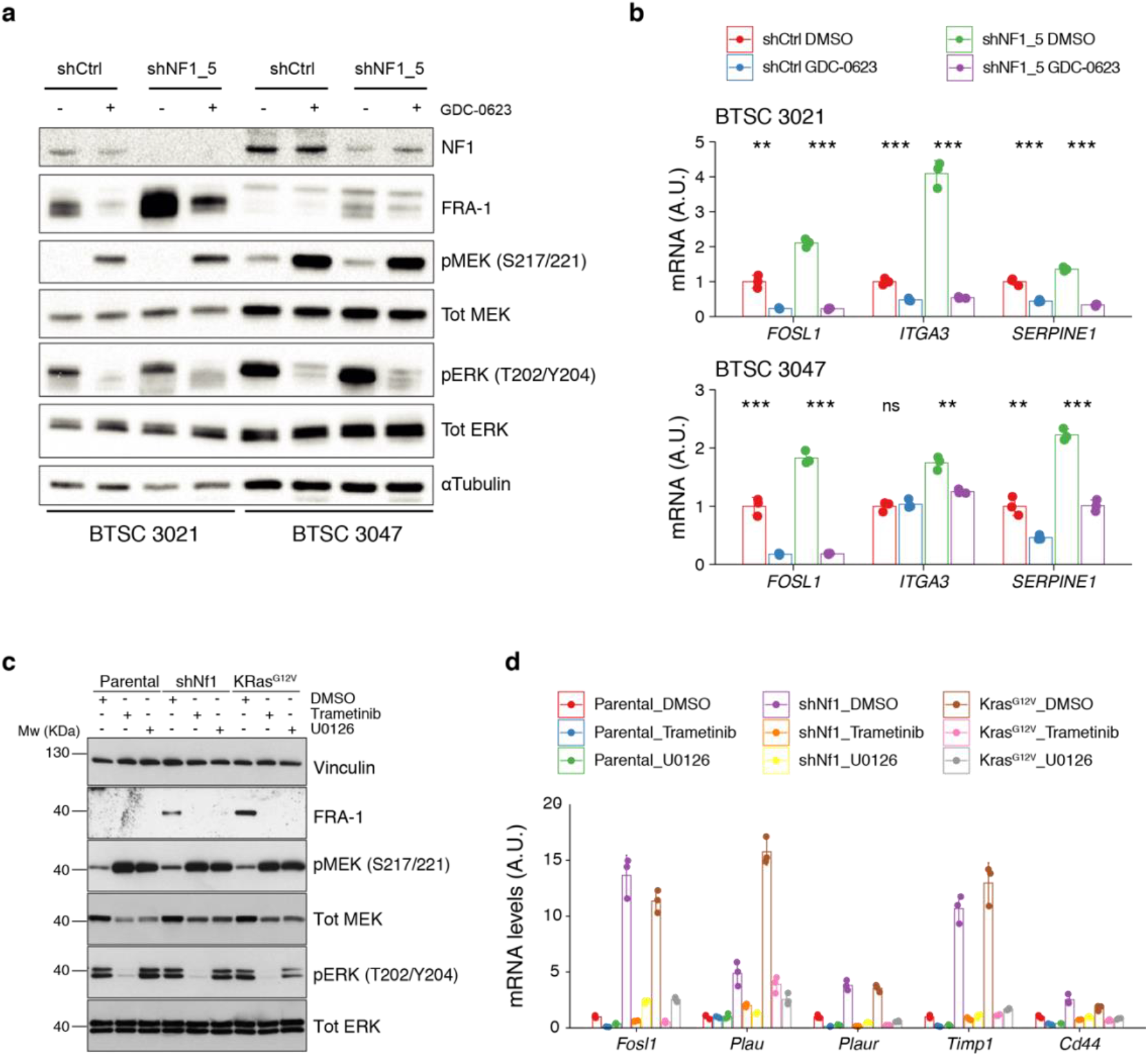
MAPK inhibition reverts the effects of NF1 silencing on *FOSL1* and mesenchymal genes expression. **a,** Western blot analysis of Non-MES cells (BTSC 3021 and 3047) transduced with shCtrl or shNF1_5 and treated with the MEK inhibitor GDC-0623 (1μM for 16 hr); α-Tubulin was used as loading control. **b,** qRT-PCR analysis of *FOSL1* and the MES genes *ITGA3* and *SERPINE1*, in samples treated as in (**a**). Data are presented as mean ± SD (n=3), normalized to 18s rRNA expression; Student’s t test of DMSO vs GDC-0623 (either shCtrl or shNF1_5), **P ≤ 0.01, ***P ≤ 0.001, ns = not-significant. **c,** Western blot analysis using the specified antibodies of p53-null NSCs, parental and infected with sh*Nf1* or *Kras^G12V^* and treated for 16 hr with the MAPK inhibitors Trametinib (200nM) or U0126 (10μM); Vinculin was used as loading control. **d,** qRT-PCR analysis of *Fosl1* and the MES genes (*Plau*, *Plaur*, *Timp1* and *Cd44*), in samples treated as in (**c**). Data are presented as mean ± SD (n=3), normalized to *Actin* expression.

**Extended Data Fig. 5.**
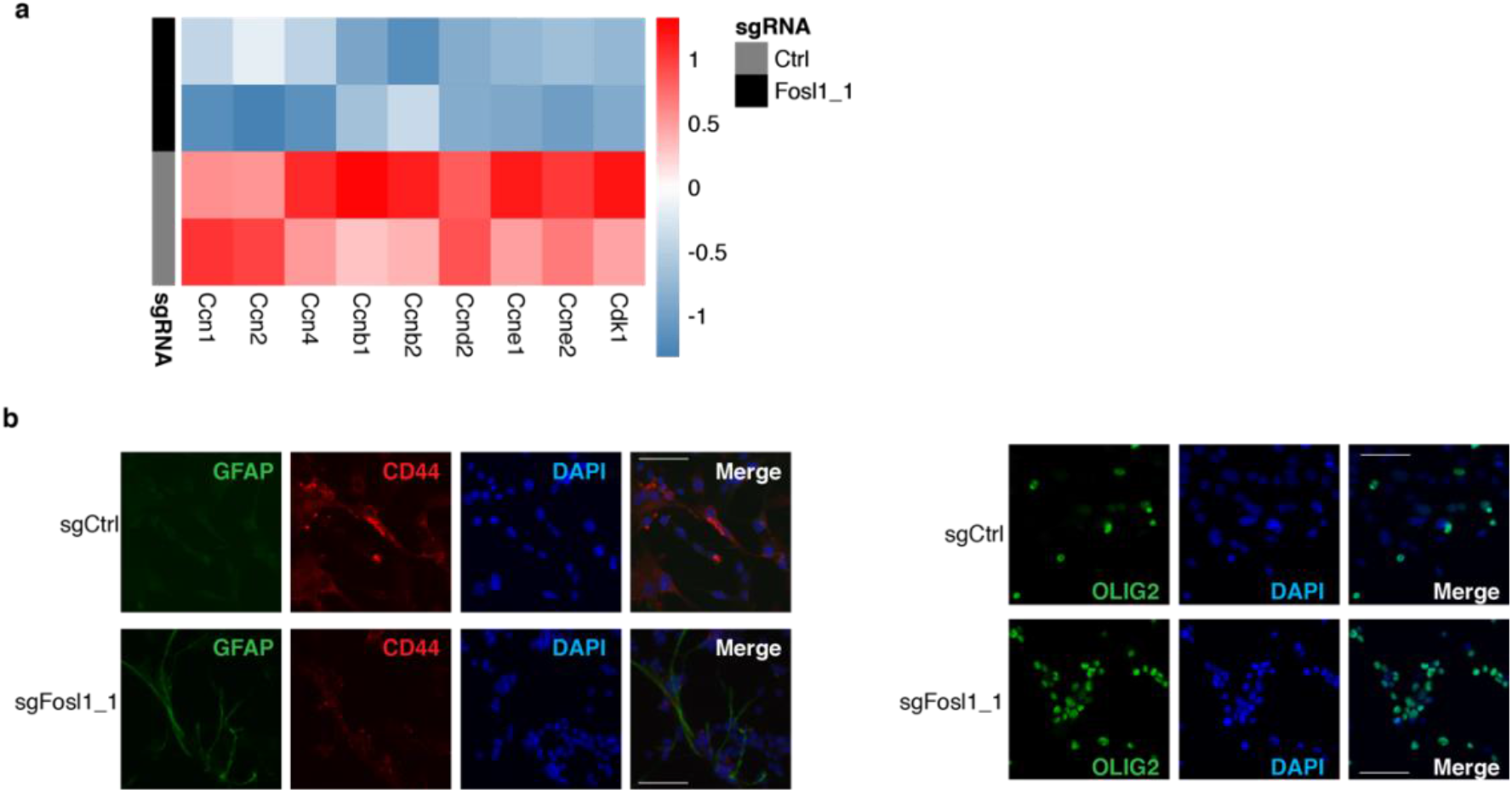
*Fosl1* loss is associated with the reduction of proliferative genes and increase in differentiation genes. **a,** Heatmap showing a reduction in expression of cell-cycle regulators in sg*Fosl1*_1, as compared to sgCtrl p53-null *Kras^G12V^* NSCs. **b,** Representative images of immunofluorescence staining of the indicated markers in sgCtrl and sg*Fosl1*_1 p53-null *Kras^G12V^* NSCs plated on laminin-coated coverslips. Scale bars represent 50 μm.

**Extended Data Fig. S6.**
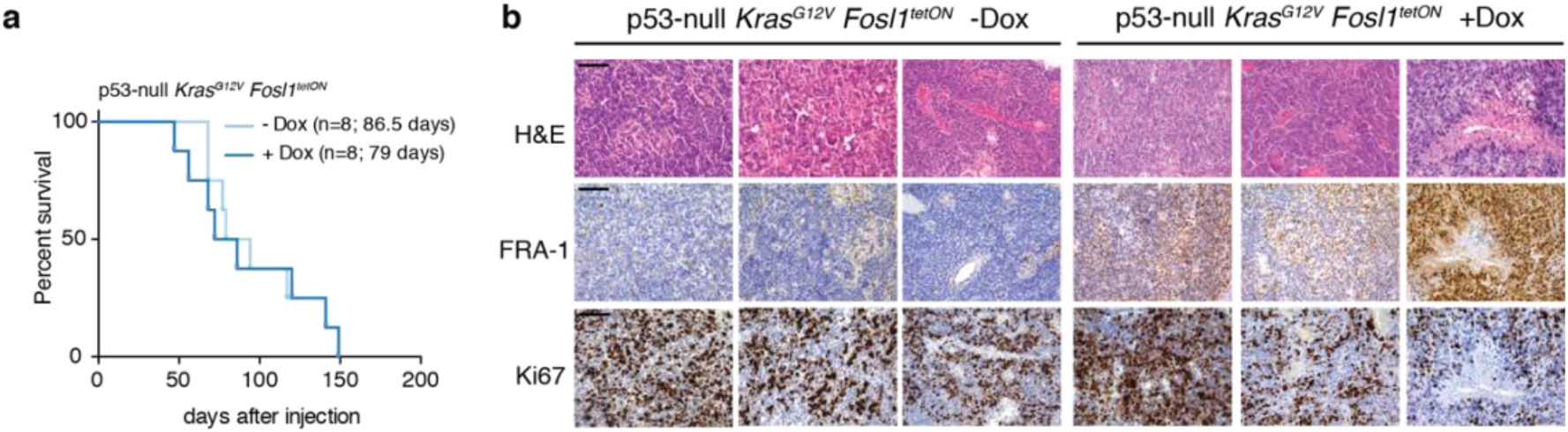
Characterization of Fosl1 overexpressing mouse tumors. **a,** Kaplan-Meier survival curves of C57BL/6J wildtype mice injected with p53-null *Kras^G12V^ Fosl1^tetON^* NSCs subjected to Dox diet (n=8) or kept as controls (n=8); Log-rank P value = 0.814. **b,** Hematoxylin and eosin (H&E) and immunohistochemical staining, using the indicated antibodies, of representative –Dox and +Dox tumors. Scale bars represent 100 μm.

**Extended Data Fig. 7.**
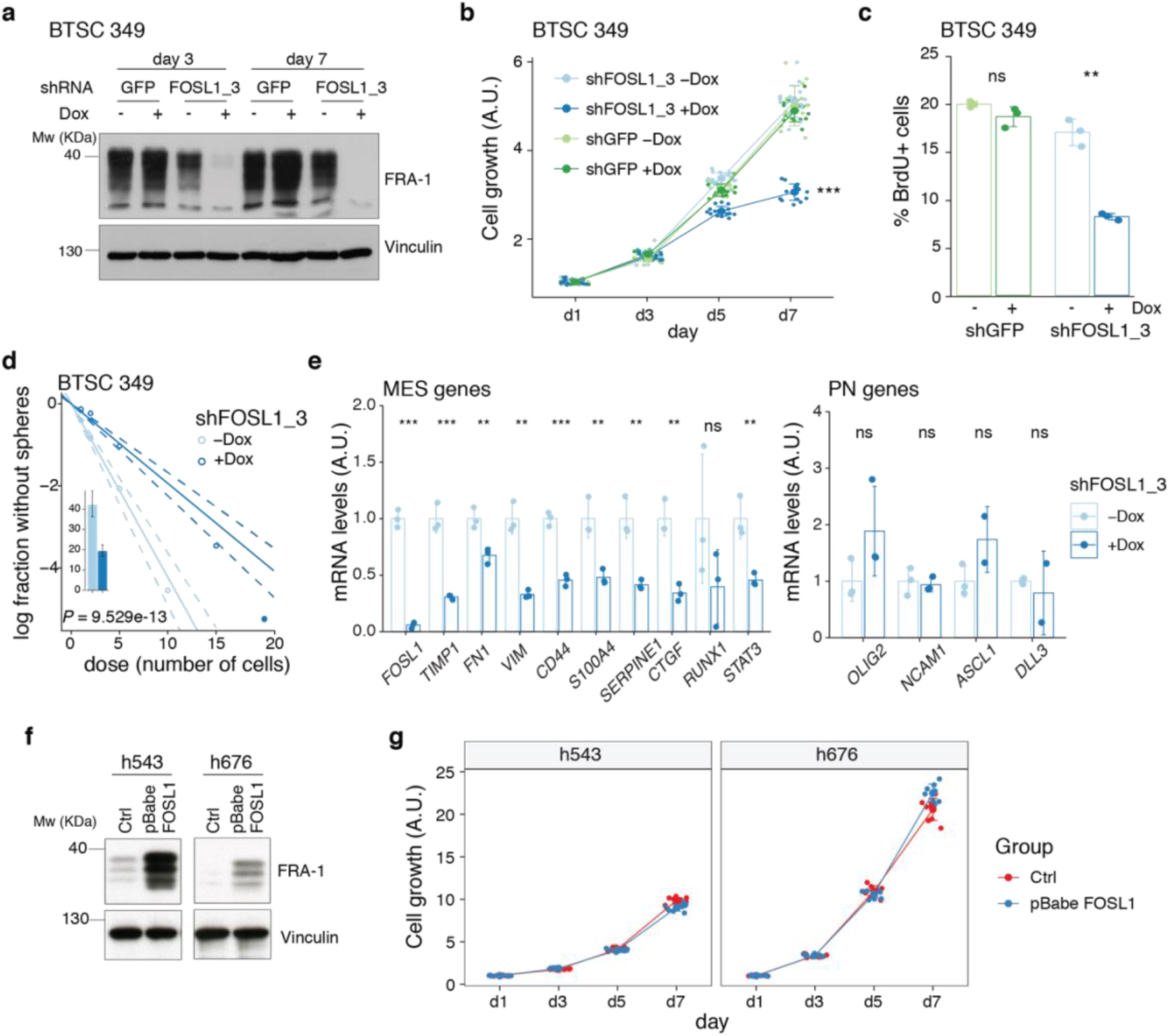
Further characterization of *FOSL1* role in human BTSCS. **a,** Western blot detection of FRA-1 in BTSC 349 upon transduction with inducible shGFP (control) or sh*FOSL1*_3, analyzed after 3 and 7 days of Dox treatment; Vinculin used as loading control. **b,** Cell growth of BTSC 349 shGFP and sh*FOSL1*_3 cells, in absence or presence of Dox, measured by MTT assay; absorbance values were normalized to day 1. Data from a representative of three independent experiments are presented as mean ± SD (n=15). Student’s t test on day 7, relative to sh*FOSL1*_3 –Dox: ***P ≤ 0.001. **c,** BrdU incorporation of BTSC 349 shGFP and sh*FOSL1*_3, in absence or presence of Dox, analyzed by flow cytometry. Data from a representative of two independent experiments are presented as mean ± SD (n=3). Student’s t test, relative to the respective control (–Dox): ns = not significant, **P ≤ 0.01. **d,** Representative limiting dilution analysis on BTSC 349 sh*FOSL1*_3 in presence or absence of Dox, calculated with extreme limiting dilution assay (ELDA) analysis; P < 0.0001. **e,** mRNA expression of *FOSL1*, MES and PN genes in BTSC 349 sh*FOSL1*_3 cells in absence or presence of Dox for 3 days. Data from a representative of three experiments are presented as mean ± SD (n=3), normalized to *GAPDH* expression. Student’s t test, relative to –Dox: ns = not-significant, *P ≤ 0.05, **P ≤ 0.01, ***P ≤ 0.001. **f,** Western blot detection of FRA-1 in h543 and h676 upon transduction with pBabe (control) or pBabe-FOSL1; Vinculin used as loading control. **g,** Cell growth of cells as in (**f**), measured by MTT assay; absorbance values were normalized to day 1. Data from a representative of three independent experiments are presented as mean ± SD.

**Extended Data Fig. 8.**
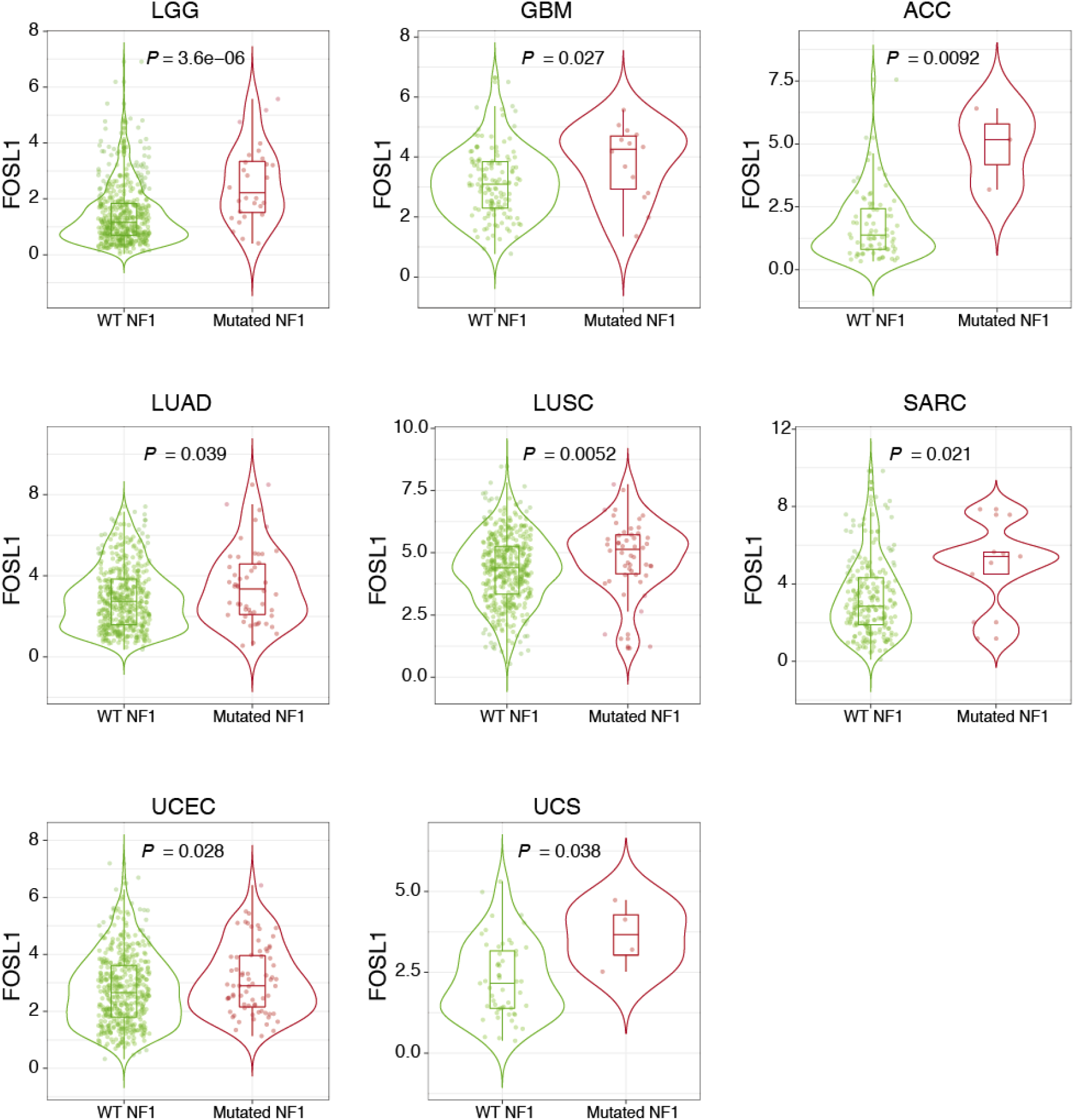
NF1 mutations are associated with higher *FOSL1* expression in multiple cancer types. *FOSL1* mRNA expression in the indicated tumors of the TCGA, stratified according to *NF1* mutational status: LGG = Low-grade glioma, GBM = Glioblastoma, ACC = Adrenocortical carcinoma, LUAD = Lung adenocarcinoma, LUSC = Lung squamous cell carcinoma, SARC = Sarcoma, UCEC = Uterine Corpus Endometrial Carcinoma, UCS = Uterine Carcinosarcoma. Wilcoxon P values are indicated.

**Extended Data Fig. 9.**
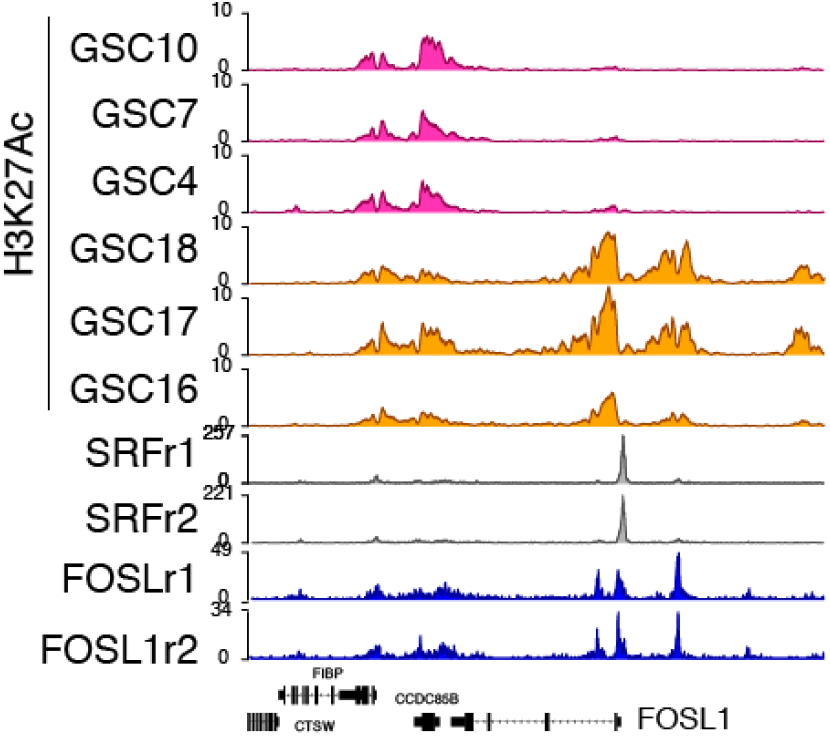
Enrichment of H3K27Ac over SRF peaks on FOSL1 promoter in MES BTSCs. SRF and FOSL1 binding sites at *FOSL1* promoter are decorated with H3K27Ac in MES cell lines.

## References

1. Louis, D. N. et al. The 2016 World Health Organization Classification of Tumors of the Central Nervous System: a summary. Acta Neuropathol. 131, 803–820 (2016).

2. Brennan, C. W. et al. The somatic genomic landscape of glioblastoma. Cell 155, 462–477 (2013).

3. Verhaak, R. G. W. et al. Integrated Genomic Analysis Identifies Clinically Relevant Subtypes of Glioblastoma Characterized by Abnormalities in PDGFRA, IDH1, EGFR, and NF1. Cancer Cell 17, 98–110 (2010).

4. The Cancer Genome Atlas Research Network. Comprehensive genomic characterization defines human glioblastoma genes and core pathways. Nature 455, 1061–1068 (2008).

5. Phillips, H. S. et al. Molecular subclasses of high-grade glioma predict prognosis, delineate a pattern of disease progression, and resemble stages in neurogenesis. Cancer Cell 9, 157–173 (2006).

6. Wang, Q. et al. Tumor Evolution of Glioma-Intrinsic Gene Expression Subtypes Associates with Immunological Changes in the Microenvironment. Cancer Cell 32, 42–56.e6 (2017).

7. Bhat, K. P. L. et al. The transcriptional coactivator TAZ regulates mesenchymal differentiation in malignant glioma. Genes Dev. 25, 2594–2609 (2011).

8. Carro, M. S. et al. The transcriptional network for mesenchymal transformation of brain tumours. Nature 463, 318–325 (2010).

9. Cooper, L. A. D. et al. The Tumor Microenvironment Strongly Impacts Master Transcriptional Regulators and Gene Expression Class of Glioblastoma. Am. J. Pathol. 180, 2108–2119 (2012).

10. Patel, A. P. et al. Single-cell RNA-seq highlights intratumoral heterogeneity in primary glioblastoma. Science 344, 1396–1401 (2014).

11. Sottoriva, A. et al. Intratumor heterogeneity in human glioblastoma reflects cancer evolutionary dynamics. Proc. Natl. Acad. Sci. 110, 4009–4014 (2013).

12. Fedele, M., Cerchia, L., Pegoraro, S., Sgarra, R. & Manfioletti, G. Proneural-Mesenchymal Transition: Phenotypic Plasticity to Acquire Multitherapy Resistance in Glioblastoma. Int. J. Mol. Sci. 20, 2746 (2019).

13. Ozawa, T. et al. Most human non-GCIMP glioblastoma subtypes evolve from a common proneural-like precursor glioma. Cancer Cell 26, 288–300 (2014).

14. Bhat, K. P. L. et al. Mesenchymal Differentiation Mediated by NF-κB Promotes Radiation Resistance in Glioblastoma. Cancer Cell 24, 331–346 (2013).

15. Halliday, J. et al. In vivo radiation response of proneural glioma characterized by protective p53 transcriptional program and proneural-mesenchymal shift. Proc. Natl. Acad. Sci. 111, 5248–5253 (2014).

16. Mani, S. A. et al. The epithelial-mesenchymal transition generates cells with properties of stem cells. Cell 133, 704–715 (2008).

17. Tam, W. L. & Weinberg, R. A. The epigenetics of epithelial-mesenchymal plasticity in cancer. Nat. Med. 19, 1438–1449 (2013).

18. Ye, X. et al. Distinct EMT programs control normal mammary stem cells and tumour-initiating cells. Nature 525, 256–260 (2015).

19. Bao, S. et al. Glioma stem cells promote radioresistance by preferential activation of the DNA damage response. Nature 444, 756–760 (2006).

20. Chen, J. et al. A restricted cell population propagates glioblastoma growth after chemotherapy. Nature 488, 522 (2012).

21. Chiappetta, G. et al. FRA-1 protein overexpression is a feature of hyperplastic and neoplastic breast disorders. BMC Cancer 7, 17 (2007).

22. Gao, X.-Q., Ge, Y.-S., Shu, Q.-H. & Ma, H.-X. Expression of Fra-1 in human hepatocellular carcinoma and its prognostic significance. Tumour Biol. 39, 1010428317709635 (2017).

23. Usui, A. et al. The molecular role of Fra-1 and its prognostic significance in human esophageal squamous cell carcinoma. Cancer 118, 3387–3396 (2012).

24. Vallejo, A. et al. An integrative approach unveils FOSL1 as an oncogene vulnerability in KRAS-driven lung and pancreatic cancer. Nat. Commun. 8, (2017).

25. Wu, J. et al. MicroRNA-195-5p, a new regulator of Fra-1, suppresses the migration and invasion of prostate cancer cells. J. Transl. Med. 13, 289 (2015).

26. Xu, H. et al. Prognostic value from integrative analysis of transcription factors c-Jun and Fra-1 in oral squamous cell carcinoma: a multicenter cohort study. Sci. Rep. 7, 7522 (2017).

27. Andreolas, C., Kalogeropoulou, M., Voulgari, A. & Pintzas, A. Fra-1 regulates vimentin during Ha-RAS-induced epithelial mesenchymal transition in human colon carcinoma cells. Int. J. cancer 122, 1745–1756 (2008).

28. Bakiri, L. et al. Fra-1/AP-1 induces EMT in mammary epithelial cells by modulating Zeb1/2 and TGFβ expression. Cell Death Differ. 22, 336–350 (2015).

29. Diesch, J. et al. Widespread FRA1-Dependent Control of Mesenchymal Transdifferentiation Programs in Colorectal Cancer Cells. PLoS One 9, 1–11 (2014).

30. Debinski, W. & Gibo, D. M. Fos-related antigen 1 modulates malignant features of glioma cells. Mol. Cancer Res. 3, 237–249 (2005).

31. Basso, K. et al. Reverse engineering of regulatory networks in human B cells. Nat. Genet. 37, 382–90 (2005).

32. Castro, M. A. A. et al. Regulators of genetic risk of breast cancer identified by integrative network analysis. Nat. Genet. 48, 12–21 (2016).

33. Fletcher, M. N. C. et al. Master regulators of FGFR2 signalling and breast cancer risk. Nat. Commun. 4, 2464 (2013).

34. Ceccarelli, M. et al. Molecular Profiling Reveals Biologically Discrete Subsets and Pathways of Progression in Diffuse Glioma. Cell 164, 550–63 (2016).

35. Zhao, Z. et al. Comprehensive RNA-seq transcriptomic profiling in the malignant progression of gliomas. Sci. Data 4, 1–7 (2017).

36. Wang, J. et al. Clonal evolution of glioblastoma under therapy. Nat. Genet. 48, 768–76 (2016).

37. McCormick, F. GAP as ras effector or negative regulator? Mol. Carcinog. 3, 185–7 (1990).

38. Ricci-Vitiani, L. et al. Mesenchymal differentiation of glioblastoma stem cells. Cell Death Differ. 15, 1491–1498 (2008).

39. Tso, C.-L. et al. Primary Glioblastomas Express Mesenchymal Stem-Like Properties. Mol. Cancer Res. 4, 607–619 (2006).

40. Oldrini, B. et al. Somatic genome editing with the RCAS-TVA-CRISPR-Cas9 system for precision tumor modeling. Nat. Commun. 9, 1466 (2018).

41. Gargiulo, G. et al. InVivo RNAi Screen for BMI1 Targets Identifies TGF-β/BMP-ER Stress Pathways as Key Regulators of Neural- and Malignant Glioma-Stem Cell Homeostasis. Cancer Cell 23, 660–676 (2013).

42. Holland, E. C. et al. Combined activation of Ras and Akt in neural progenitors induces glioblastoma formation in mice. Nat. Genet. 25, 55–57 (2000).

43. Uhrbom, L. et al. Ink4a-Arf loss cooperates with KRas activation in astrocytes and neural progenitors to generate glioblastomas of various morphologies depending on activated Akt. Cancer Res. 62, 5551–5558 (2002).

44. Friedmann-Morvinski, D. et al. Dedifferentiation of neurons and astrocytes by oncogenes can induce gliomas in mice. Science 338, 1080–1084 (2012).

45. Koschmann, C. et al. ATRX loss promotes tumor growth and impairs nonhomologous end joining DNA repair in glioma. Sci. Transl. Med. 8, 328ra28 (2016).

46. Muñoz, D. M. et al. Loss of p53 cooperates with K-ras activation to induce glioma formation in a region-independent manner. Glia 61, 1862–1872 (2013).

47. Belteki, G. et al. Conditional and inducible transgene expression in mice through the combinatorial use of Cre-mediated recombination and tetracycline induction. Nucleic Acids Res. 33, 1–10 (2005).

48. Hasenfuss, S. C., Bakiri, L., Thomsen, M. K., Hamacher, R. & Wagner, E. F. Activator protein 1 transcription factor fos-related antigen 1 (fra-1) is dispensable for murine liver fibrosis, but modulates xenobiotic metabolism. Hepatology 59, 261–273 (2014).

49. Mack, S. C. et al. Chromatin landscapes reveal developmentally encoded transcriptional states that define human glioblastoma. J. Exp. Med. 216, 1071–1090 (2019).

50. Neftel, C. et al. An Integrative Model of Cellular States, Plasticity, and Genetics for Glioblastoma. Cell 178, 835–849.e21 (2019).

51. Wang, L. et al. The phenotypes of proliferating glioblastoma cells reside on a single axis of variation. Cancer Discov. CD-19-0329 (2019). doi:10.1158/2159-8290.CD-19-0329

52. Liu, H. et al. Aberrantly expressed Fra-1 by IL-6/STAT3 transactivation promotes colorectal cancer aggressiveness through epithelial-mesenchymal transition. Carcinogenesis 36, 459–468 (2015).

53. Casalino, L., De Cesare, D. & Verde, P. Accumulation of Fra-1 in ras-Transformed Cells Depends on Both Transcriptional Autoregulation and MEK-Dependent Posttranslational Stabilization. Mol. Cell. Biol. 23, 4401–4415 (2003).

54. Verde, P., Casalino, L., Talotta, F., Yaniv, M. & Weitzman, J. B. Deciphering AP-1 function in tumorigenesis: Fra-ternizing on target promoters. Cell Cycle 6, 2633–2639 (2007).

55. Esnault, C. et al. ERK-Induced Activation of TCF Family of SRF Cofactors Initiates a Chromatin Modification Cascade Associated with Transcription. Mol. Cell 65, 1081–1095.e5 (2017).

56. Lau, E. Y. T. et al. Cancer-Associated Fibroblasts Regulate Tumor-Initiating Cell Plasticity in Hepatocellular Carcinoma through c-Met/FRA1/HEY1 Signaling. Cell Rep. 15, 1175–1189 (2016).

57. Eferl, R. & Wagner, E. F. AP-1: A double-edged sword in tumorigenesis. Nat. Rev. Cancer 3, 859–868 (2003).

58. Cusulin, C. et al. Precursor States of Brain Tumor Initiating Cell Lines Are Predictive of Survival in Xenografts and Associated with Glioblastoma Subtypes. Stem Cell Reports 5, 1–9 (2015).

59. Mao, P. et al. Mesenchymal glioma stem cells are maintained by activated glycolytic metabolism involving aldehyde dehydrogenase 1A3. Proc. Natl. Acad. Sci. U. S. A. 110, 8644–9 (2013).

60. Günther, H. S. et al. Glioblastoma-derived stem cell-enriched cultures form distinct subgroups according to molecular and phenotypic criteria. Oncogene 27, 2897–909 (2008).

61. Graña*, O., Rubio-Camarillo, M., Fdez-Riverola, F. & Glez-Peña, D.G. P. and Nextpresso D. Next Generation Sequencing Expression Analysis Pipeline. Current Bioinformatics 13, 583–591 (2018).

62. Gautier, L., Cope, L., Bolstad, B. M. & Irizarry, R. A. affy-analysis of Affymetrix GeneChip data at the probe level. BIOINFORMATICS 20, 307–315 (2004).

63. Ritchie, M. E. et al. limma powers differential expression analyses for RNA-sequencing and microarray studies. Nucleic Acids Res. 43, (2015).

64. Li, T. et al. TIMER2.0 for analysis of tumor-infiltrating immune cells. Nucleic Acids Res. 48, 509–514 (2020).

65. Bowman, R. L., Wang, Q., Carro, A., Verhaak, R. G. W. & Squatrito, M. GlioVis data portal for visualization and analysis of brain tumor expression datasets. Neuro. Oncol. 19, 139–141 (2017).

66. Hennig, B. P. et al. Large-scale low-cost NGS library preparation using a robust Tn5 purification and tagmentation protocol. G3 Genes, Genomes, Genet. 8, 79–89 (2018).

67. Buenrostro, J. D., Giresi, P. G., Zaba, L. C., Chang, H. Y. & Greenleaf, W. J. Transposition of native chromatin for fast and sensitive epigenomic profiling of open chromatin, DNA-binding proteins and nucleosome position. Nat. Methods 10, 1213–1218 (2013).

68. Langmead, B. & Salzberg, S. L. Fast gapped-read alignment with Bowtie 2. Nat. Methods 9, 357–359 (2012).

69. Schep, A. N., Wu, B., Buenrostro, J. D. & Greenleaf, W. J. ChromVAR: Inferring transcription-factor-associated accessibility from single-cell epigenomic data. Nat. Methods 14, 975–978 (2017).

70. Yu, G., Wang, L. G., Han, Y. & He, Q. Y. ClusterProfiler: An R package for comparing biological themes among gene clusters. Omi. A J. Integr. Biol. 16, 284–287 (2012).

71. Ramírez, F. et al. deepTools2: a next generation web server for deep-sequencing data analysis. Web Serv. issue Publ. online 44, (2016).

72. Gel, B. & Serra, E. karyoploteR: an R/Bioconductor package to plot customizable genomes displaying arbitrary data and 2 CIBERONC. doi:10.1093/bioinformatics/btx346

73. Zhang, Y. et al. Model-based analysis of ChIP-Seq (MACS). Genome Biol. 9, R137 (2008).

74. Suvà, M. L. et al. Reconstructing and reprogramming the tumor-propagating potential of glioblastoma stem-like cells. Cell 157, 580–594 (2014).

75. Love, M. I., Huber, W. & Anders, S. Moderated estimation of fold change and dispersion for RNA-seq data with DESeq2. Genome Biol. 15, 550 (2014).

76. Younesy, H. et al. ChAsE: chromatin analysis and exploration tool. Bioinformatics 32, 3324–3326 (2016).

77. Matsuo, K. et al. Fos/1 is a transcriptional target of c-Fos during osteoclast differentiation. Nat. Genet. 24, 184–187 (2000).

78. Annibali, D. et al. Myc inhibition is effective against glioma and reveals a role for Myc in proficient mitosis. Nat. Commun. 5, 1–11 (2014).

79. Mansuy, I. M. & Bujard, H. Tetracycline-regulated gene expression in the brain. Curr. Opin. Neurobiol. 10, 593–596 (2000).

80. Hambardzumyan, D., Amankulor, N. M., Helmy, K. Y., Becher, O. J. & Holland, E. C. Modeling Adult Gliomas Using RCAS/t-va Technology. Transl. Oncol. 2, 89–95 (2009).

81. Dobin, A. et al. STAR: ultrafast universal RNA-seq aligner. Bioinformatics 29, 15–21 (2013).

82. Anders, S., Pyl, P. T. & Huber, W. HTSeq--a Python framework to work with high-throughput sequencing data. Bioinformatics 31, 166–9 (2015).

83. Sandberg, C. J. et al. Comparison of glioma stem cells to neural stem cells from the adult human brain identifies dysregulated Wnt-signaling and a fingerprint associated with clinical outcome. Exp. Cell Res. 319, 2230–2243 (2013).

84. Bazzoli, E. et al. MEF Promotes Stemness in the Pathogenesis of Gliomas. Cell Stem Cell 11, 836–844 (2012).

85. Subramanian, A. et al. Gene set enrichment analysis: a knowledge-based approach for interpreting genome-wide expression profiles. Proc. Natl. Acad. Sci. U. S. A. 102, 15545–15550 (2005).

